# Impaired stem cell migration and divisions in Duchenne Muscular Dystrophy revealed by live imaging

**DOI:** 10.1101/2025.03.13.643016

**Authors:** Liza Sarde, Gaëlle Letort, Hugo Varet, Vincent Laville, Julien Fernandes, Shahragim Tajbakhsh, Brendan Evano

**Affiliations:** Stem Cells and Development, Department of Developmental and Stem Cell Biology, Institut Pasteur, Université Paris Cité, 75015 Paris, France; CNRS UMR 3738, Institut Pasteur, 75015 Paris, France; Sorbonne Université, Complexité du Vivant, F-75005 Paris, France; Department of Developmental and Stem Cell Biology, Institut Pasteur, Université Paris Cité, CNRS UMR 3738, 75015 Paris, France; Bioinformatics and Biostatistics Hub, Research and Resource Centre for Scientific Informatics, Institut Pasteur, 75015 Paris, France; Institut Pasteur, Université Paris Cité, Photonic Bio-Imaging Unit, Centre de Ressources et Recherches Technologiques (UTechS-PBI, C2RT), F-75015 Paris, France

## Abstract

Dysregulation of stem cell properties is a hallmark of many pathologies, but the dynamic behaviour of stem cells in their microenvironment during disease progression remains poorly understood. Using the *mdx* mouse model of Duchenne Muscular Dystrophy, we developed innovative live-imaging of muscle stem cells (MuSCs) *in vivo*, and *ex vivo* on isolated myofibres. We show that *mdx* MuSCs have impaired migration and precocious differentiation through unbalanced symmetric divisions, driven by p38 and PI3K signalling pathways, in contrast to the p38-only dependence of healthy MuSCs. Cross-grafting shows that MuSC fate decisions are governed by intrinsic cues, whereas their migration behaviour is determined by the extracellular niche. This study provides the first dynamic analysis of dystrophic MuSC properties *in vivo*, reconciling conflicting reports on their function. Our findings establish DMD as a MuSC disease with intrinsic defects and niche dysfunction, offering strategies to restore stem cell functions for improved muscle regeneration.

## Introduction

Duchenne Muscular Dystrophy (DMD) is a severe X-linked genetic disease characterised by muscle wasting, physical incapacitation, and death around the age of 20-30. Affecting ∼1/5000 male births, DMD arises from mutations in the DMD gene encoding *Dystrophin*, a structural protein essential for myofibre integrity. *Dystrophin* deficiency leads to chronic muscle degeneration, inflammation, fibrosis, and impaired regeneration^1^. While *ex vivo* models suggest that muscle stem cell (MuSC) dysfunction contributes to DMD pathology^2^, *in vivo* dynamics of MuSCs and early myogenic cell perturbations remain unexplored.

MuSCs reside in a niche between the myofibre and basement membrane, maintaining quiescence during homeostasis. Upon activation (*e.g.* muscle injury or isolation), MuSCs proliferate and differentiate to generate or repair muscle fibres, with a subset self-renewing. Myogenic progression involves temporal expression of transcription factors PAX7 (stem), MYF5 and MYOD (commitment), and MYOGENIN (MYOG; differentiation). Symmetric and asymmetric cell divisions (SCD/ACD) regulate this process^3^. Impaired ACD has been implicated in *mdx* mice (DMD mouse model), with MuSC hyperplasia and deficient progenitor generation^4^. Although SCDs are predominant in regenerating muscles^5^, their role in normal and *mdx* models remains unexplored. Additionally, conflicting studies reported increased^6–8^ or decreased^4,9^ differentiation of dystrophic myogenic cells, leaving the balance of proliferation and differentiation through ACDs and SCDs in *mdx* MuSCs unresolved.

MuSC differentiation integrates pathways such as p38 mitogen-activated protein kinases (MAPK)^10^ and PI3K signalling^11^. p38α/ß inhibitors (SB203580) or *Mapk14* deletion (p38α-encoding gene) reduced inflammation and prevented myofibre death in *mdx* mice^12^, however, the role of p38 in *mdx* MuSC differentiation remains to be explored. Furthermore, increased Akt activation was reported in *mdx* mouse muscles and in DMD patients^13,14^, yet elevated PI3K inhibitor PTEN was reported in golden retriever muscular dystrophy (GRMD) dogs and *mdx* mouse myoblasts^15,16^. Targeting PI3K/Akt signalling may mitigate DMD progression^17^, but its role in *mdx* MuSC differentiation is poorly studied.

Cell-based therapeutic strategies have been explored for muscle wasting diseases including DMD, however, clinical trials failed, in part due to limited cell migration (reviewed in^18,19^). MuSCs are immobile during homeostasis *in vivo*^20^, and they migrate along damaged fibres during regeneration following Erk signalling activation and interaction with stromal cells^21–23^. While *in vitro* studies suggest reduced chemotaxis and impaired migration^6^, *in vivo* migration and positioning of dystrophic MuSCs remain unstudied^24^.

MuSC properties arise from complex interactions between intrinsic and extrinsic cues^20^, as evidenced by transcriptome and epigenome reprogramming after transplantation to different anatomical or age-specific environments^25^. For dystrophy therapies, tissue dispersion of engrafted myogenic cells has been enhanced by boosting intrinsic migration capacity and modulating the environment^18^. Understanding these cues is essential for developing effective therapeutic strategies for healthy and dystrophic MuSCs.

We developed novel quantitative pipelines to track MuSC migration, proliferation and differentiation *in vivo* and *ex vivo* on their myofibre niche, in healthy and *mdx* contexts. *mdx* MuSCs showed impaired migration kinetics, increased symmetric divisions towards terminal differentiation, and reduced symmetric stem cell renewal. Transplant experiments revealed that fate decisions are primarily MuSC-intrinsic while migration is largely influenced by myofibre-derived cues, indicating that these processes are uncoupled in *mdx* mice. Further, we identified differential roles for p38 MAPK and PI3K signalling in normal and dystrophic differentiation.

## Results

### Precocious differentiation and impaired migration of dystrophic MuSCs revealed by intravital imaging

Intravital imaging was used to investigate myogenic cell properties during muscle regeneration *in vivo* in wild-type (WT) and *mdx* mice. To do so, MuSCs and their descendants were genetically labelled with a membrane-GFP (mGFP) using inducible *Pax7^CreERT2^* ^26^ and *R26^mTmG^* ^27^ reporters in adult *Dmd^+/Y^* (WT) and *Dmd^mdx-βGeo/Y^* ^28^ (*mdx*) mice (Fig. 1a). The hindlimb *Flexor digitorum brevis* (FDB) muscle was imaged intravitally by two-photon microscopy due to its accessibility (Fig. 1b and Fig. S1). Of note, *mdx* fibres exhibited centrally located nuclei (46.9%, vs 3.2% in WT) (Fig. S2b-c) in resting FDBs, a sign of regeneration resulting from the chronic degeneration/regeneration as expected in DMD. To circumvent the non-synchronous nature of myofibre damage and repair in *mdx* mice, we induced acute muscle injury in the FDB muscle by injecting cardiotoxin, a snake venom, to trigger MuSC activation. Myogenic cells were visualised at 3 days post-injury (dpi) when they undergo active divisions and migration^22,23^. WT myogenic cells were mononucleated, showed an elongated/mesenchymal-like morphology with few protrusions, actively proliferated and migrated, and had a uni- or bi-polar morphology (Fig. 1c, Movie S1 and S2), as reported^29–31^. In contrast, *mdx* mice showed the presence of round and immobile mononucleated cells and multinucleated mGFP+ myotubes (Fig. 1d-e, Movie S1 and S2), suggesting faster differentiation and fusion kinetics compared with the WT. Next, we quantified migration speed and directionality (reflected by straightness and turning angle) (Fig. 1f-i). WT myogenic cells migrated at 28.5 μm/h and exhibited a straight trajectory (mean straightness = 0.75)(Fig. 1g-h), indicating active migration along the longitudinal axis of damaged muscle fibres as reported previously^22,23^. In contrast, *mdx* MuSCs showed overall reduced speed (26.2 μm/h, p = 6.1e-05), and reduced directionality (mean straightness = 0.70, p = 1.13e-08 and turning angle, mean *mdx* = 47.7° vs mean WT = 38.1°, p = 7.8e-16) (Fig. 1g-i), and the presence of round and immobile cells (Fig. 1d, Movie S2). After mitosis and completion of cytokinesis, WT cells separated rapidly from each other and migrated in opposing directions (> 95%, 30-60 min after mitosis), whereas a fraction *mdx* cells migrated in the same direction shortly after mitosis (∼20%, 30-60 min after mitosis) (Fig. 1j-l).

**Figure 1.**
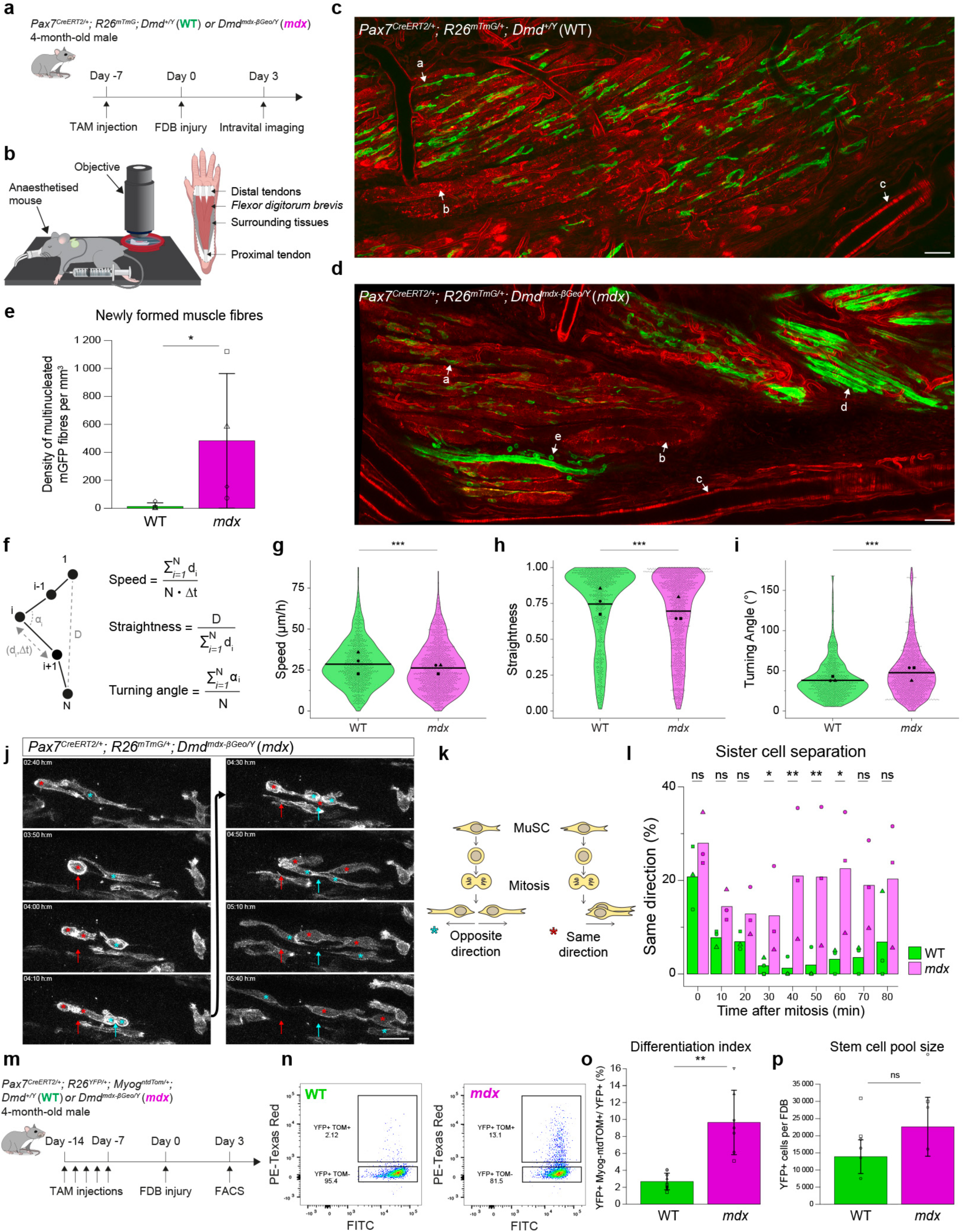
Precocious differentiation and impaired migration of dystrophic MuSCs revealed by intravital imaging. **(a)** Experimental scheme. Adult *Pax7^CreERT2/+^; R26^mTmG/+^;* WT and *mdx* 4-month-old male mice were treated with tamoxifen. The FDB was injured with cardiotoxin and imaged by intravital imaging at 3 dpi for 8-10h. **(b)** Scheme of intravital imaging. The mouse was anaesthetised with isofluorane, kept on a heat pad and hydrated every hour with NaCl 0.9% *via* a catheter. A glass coverslip was attached to the FDB muscle for imaging. See also Fig. S1. **(c-d)** Representative image of intravital imaging of injured FDBs at 3 dpi of *Pax7^CreERT2/+^; R26^mTmG/+^;* (**c**) *Dmd^+/Y^* (WT) and (**d**) *Dmd^mdx-βGeo/Y^* (*mdx)* adult mice. MuSCs and their progeny (myoblasts and newly formed myofibres) were labelled with mGFP, and the surrounding tissue was labelled with mTOMATO. Arrows: a, MuSC; b, injured FDB fibre; c, blood vessel; d, newly formed myofibre; e, static myoblast. Scale bar, 50 μm. See Movie S1 and S2. **(e)** Density of multinucleated mGFP fibres per imaging volume (mm^3^) in WT and *mdx* mice at 3 dpi. N = 4. p = 0.0265. **(f)** Calculation of motility parameters: speed, straightness and turning angle. d_i_: distance between time frames i and i+1; D: net distance between start (1) and end (N) points; Δt: time interval between consecutive time frames, α_i_: {0:180}, angle of migration path between time frames i-1, i and i+1. **(g)** Migration speed of mGFP-labelled WT and *mdx* MuSCs *in vivo* at 3 dpi. N = 3 experiments, 50-200 cells analysed/experiment/mouse. p = 6.073e-05. **(h)** Migration straightness of mGFP-labelled WT and *mdx* MuSCs *in vivo* at 3 dpi. N = 3 experiments, 50-200 cells analysed/experiment/mouse. p = 1.126e-08. **(i)** Measure of angles between consecutive time frames of mGFP-labelled WT and *mdx* MuSCs *in vivo* at 3 dpi. N = 3 experiments, 50-200 cells analysed/experiment/mouse. p = 7.819e-16. **(j)** Representative intravital imaging of myoblast migration following mitosis from *Pax7^CreERT2/+^; R26^mTmG/+^; Dmd^mdx-βGeo/Y^* (*mdx)* 4-month-old adult mouse at 3 dpi. After division, sister cells can migrate in opposite (cyan stars) or in the same direction (red stars) with respect to mitosis site (arrow). Scale bar, 20 μm. **(k)** Scheme of sister cell separation after mitosis, in same or opposite directions. **(l)** Percentage of divisions with sister cells migrating in the same direction following mitosis in WT and *mdx* mice *in vivo* at 3 dpi. N = 3 experiments. **(m)** Experimental scheme. *Pax7^CreERT2/+^; R26^YFP/+^; Myog^ntdTom/+^; Dmd^+/Y^* (WT) and *Dmd^mdx-βGeo/Y^* (*mdx)* 4-month-old male mice, treated 5X with tamoxifen. FDBs were injured with cardiotoxin and dissociated at 3 dpi to measure differentiation index (% YFP-positive, Myog-ntdTOM-positive myogenic cells) and absolute numbers of YFP-labelled myogenic cells by FACS. **(n)** Representative FACS plot of Myog-ntdTOM expression in WT (left) and *mdx* (right) YFP-labelled myogenic cells at 3 dpi. **(o)** Differentiation index of WT and *mdx* YFP-labelled myogenic cells at 3 dpi. n = 9 WT mice, n = 7 *mdx* mice, p = 0.00102. **(p)** Absolute number of WT and *mdx* YFP-labelled myogenic cells per FDB at 3 dpi. n = 6 WT mice, n = 4 *mdx* mice. p = 0.257. Statistical tests: **(g-i, l)** Linear mixed models; **(e, o-p)** Wilcoxon test. Horizontal lines in violin plot represent the mean. * *p* < 0.05, ** *p* < 0.01, *** *p* < 0.001.

To investigate further the differentiation dynamics of MuSCs, we used a *Myog^ntdTom^* ^32^ knock-in reporter for differentiation together with *Pax7^CreERT2^* and *R26^YFP^* ^33^ alleles for labelling MuSCs and their progeny (Fig. 1m). Cytometry analyses at 3 dpi showed a higher differentiation index of *mdx* MuSCs compared with WT (9.6% vs 2.7% respectively, p = 1.0e-03, Fig. 1n-o), but no difference in absolute numbers of mononucleated myogenic cells (Fig. 1p).

Altogether, our observations point to precocious differentiation and impaired migration kinetics of *mdx* MuSCs *in vivo* during muscle regeneration. To further explore the cellular and molecular mechanisms underlying these defects, we developed an *ex vivo* model designed to: *i)* recapitulate *in vivo* phenotypes, *ii)* enable dynamic tracking of migration and fate decisions, and *iii)* facilitate perturbation studies.

### Impaired symmetric divisions, differentiation and migration kinetics of dystrophic MuSCs in their myofibre niche

MuSC fate decisions have been extensively studied using *ex vivo* cultures of MuSCs on primary myofibres in suspension, a system that partially preserves the MuSC niche^20^, followed by static analysis of sister cell fates^4,34–37^. However, MuSCs are highly motile on cultured myofibres^38^, thereby creating uncertainty in identifying daughter cells by static imaging. To overcome these limitations, we designed a confinement system with microwells that enables suspension culture, live imaging and retrospective immunostaining of single myofibres (Fig. 2a-b and Fig. S2a). FDB myofibres were selected due to their small size (length ∼ 700 μm), which facilitates their manipulation and full-length imaging. FDB fibres from adult *Pax7^CreERT2^*; *R26^mTmG^* mice in microwells were filmed for 72h to monitor MuSC activation kinetics (Fig. 2c and Movie S3). MuSCs transiting from quiescence to early activation were mostly immobile and showed dynamic cellular protrusions, as previously reported^30,31,39^. They migrated more rapidly, proliferated, and underwent cell divisions where daughter cells were tracked continuously for up to five consecutive divisions (Movie S3). Therefore, our culture and imaging system allowed quantitative assessment of multiple parameters from quiescence to differentiation on the myofibre niche.

**Figure 2.**
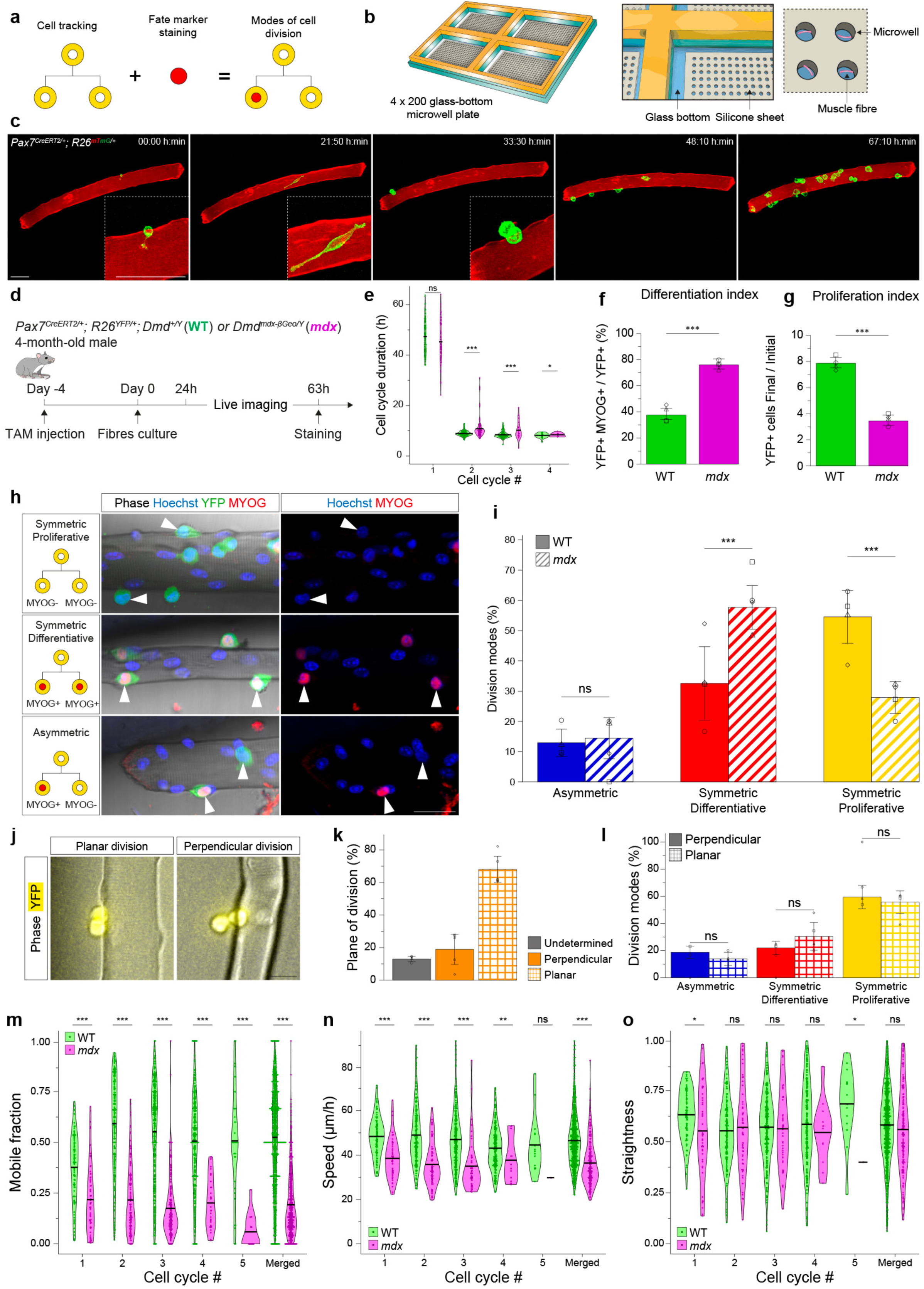
Impaired symmetric divisions, differentiation and migration kinetics of dystrophic MuSCs in their myofibre niche. **(a)** Design of an *ex vivo* live-imaging pipeline to record activation timing, cell cycle duration, proliferation and differentiation rates, modes of division (symmetric vs asymmetric, live or with end point immunostaining) and migration parameters (speed, straightness, distance) of WT and *mdx* MuSCs in their myofibre niche. **(b)** Scheme of microwell plate. A 4-well Plexiglass support, glass coverslips and silicone microwells are sealed together to allow single FDB myofibres culture in suspension, live imaging and endpoint immunostaining. **(c)** Snapshots of a representative example of live imaging of FDB fibres in microwells. An adult *Pax7^CreERT2/+^; R26^mTmG/+^* mouse was treated with tamoxifen, FDB fibres were isolated in microwells and imaged for 3 days. Scale bar, 50 μm. See Movie S3. **(d)** Experimental scheme. *Pax7^CreERT2/+^; R26^YFP/+^; Dmd^+/Y^* (WT) and *Dmd^mdx-βGeo/Y^* (*mdx)* WT and *mdx* 4-month-old male mice were treated once with tamoxifen. FDB fibres and associated MuSCs were isolated, cultured and filmed in microwells for ∼ 63h, fixed and immunostained. **(e)** Cell cycle duration (h) for all cell cycle for *Pax7^CreERT2/+^; R26^YFP/+^*; WT and *mdx* mice. N = 4 experiments. **(f)** Differentiation index of WT and *mdx* YFP-labelled myogenic cells (∼ 63h of culture). N = 4 experiments. p = 4.308e-06. **(g)** Proliferation index of WT and *mdx* YFP-labelled myogenic cells (∼ 63h of culture). N = 4 experiments. p = 8.875e-11. **(h)** Representative examples of symmetric proliferative (top), symmetric differentiative (middle) and asymmetric divisions (bottom) of YFP-labelled myogenic cells. Scale bar, 30 μm. **(i)** Modes of cell divisions of WT and *mdx* YFP-labelled myogenic cells. N = 4 experiments. p-values: ACD (WT vs *mdx*) = 0.77; SCDd (WT vs *mdx*) = 3.3e-05; SCDp (WT vs *mdx*) = 1.2e-05. **(j)** Representative examples of divisions of YFP-labelled myogenic cells planar (left) or perpendicular (right) along the long axis of the myofibre. Scale bar, 20 μm. **(k)** Percentage of planar and perpendicular divisions of YFP-labelled myogenic cells. N = 4 experiments. The orientation of 13% of divisions could not be determined due to their dynamic nature and lack of spatiotemporal resolution with our imaging modality (see Methods). **(l)** Modes of cell divisions of WT YFP-labelled myogenic cells following perpendicular or planar divisions. N = 4 experiments. p-values: ACD (planar vs perpendicular) = 0.5; SCDd (planar vs perpendicular) = 0.35; SCDp (planar vs perpendicular) = 0.71. **(m)** Mobile fraction of WT and *mdx* YFP-labelled myogenic cells, for individual and merged cell cycles. N = 4 experiments. **(n)** Migration speed of the mobile fraction of WT and *mdx* YFP-labelled myogenic cells, for individual and merged cell cycles. N = 4 experiments. **(o)** Migration straightness of the mobile fraction of WT and *mdx* YFP-labelled myogenic cells, for individual and merged cell cycles. N = 4 experiments. Statistical tests: **(f-g)** Wilcoxon test; **(e, i, l-o)** Linear mixed models. Horizontal lines in violin plots represent the mean. * *p* < 0.05, ** *p* < 0.01, *** *p* < 0.001.

To assess the properties of dystrophic myogenic cells, we isolated FDB fibres from 4-month-old *Pax7^CreERT2^*; *R26^YFP^* WT or *mdx* mice in microwells, tracked YFP-labelled MuSCs and their progeny, recorded cell migration, and immunostained for MYOG to reconstruct the lineage and modes of cell divisions retrospectively (Fig. 2a,d). The initial number of MuSCs per fibre was similar between WT and *mdx* mice (mean WT = 1.8, mean *mdx* = 2.3, p = 0.055, Fig. S2d). Further, sampling of the MuSC population by partial recombination of the *R26^YFP^* allele through *Pax7^CreERT2^* and low-dose tamoxifen (see Methods) yielded similar results between WT and *mdx* (mean recombination WT = 57.4%, *mdx* = 60.6%, p = 0.88, Fig. S2e), indicating no overt bias due to heterogeneities in behaviour among clonally labelled WT and *mdx* MuSCs. As previously reported^38,40^, WT MuSCs executed their first division on average around 47.3h, then divided approximately every 8.5h (Fig. 2e), further validating the fidelity of our culture system. Of note, daughter cells took about 2.3h to fully disconnect post-mitosis (Fig. S2f), highlighting this time window for accurate identification of sister cells and their fate (a)symmetry^4,34–37^. Further, except for the first cell cycle where there was no difference in cell cycle kinetics, *mdx* MuSCs divided more slowly than WT cells (Fig. 2e).

Importantly, *mdx* MuSCs showed increased differentiation (*mdx* = 76%, WT = 38%, p = 4.31e-06, Fig. 2f) and reduced proliferation (*mdx* = 3.5, WT = 7.9, p = 8.88e-11, Fig. 2g) indexes compared to WT, thereby confirming our *in vivo* observations (Fig. 1). Further, we observed symmetric proliferative (SCDp; sister cells MYOG-, Fig. 2h top) and differentiative (SCDd, sister cells MYOG+, Fig. 2h middle) divisions, and asymmetric cell divisions (one sister MYOG+, Fig. 2h bottom), as previously reported^5,40,41^. WT MuSCs divided mostly symmetrically (ACD = 12.9%; SCDd = 32.5%, SCDp = 54.5%, Fig. 2i), as *in vivo*^5^. Unexpectedly, *mdx* MuSCs showed no impairment of ACD^4^ (14.4%, p-value vs WT = 0.77) but drastic alterations of symmetric divisions, with increased SCDd (57.7%, p-value vs WT = 3.3e-05) and decreased SCDp (27.9%, p-value vs WT = 1.2e-05) (Fig. 2i). Further, the mode of cell division was reported to be impacted by the extrinsic microenvironment and the orientation of the mitotic spindle apparatus (see^42^). To explore this possibility, we determined if the orientation of cell division relative to the myofibre was related to the modes of cell divisions, as early ACD decisions were reported to result from differential exposure of sister cells to basal lamina and myofibre-derived cues^4,34–37^. We observed planar (68.0%, Fig. 2j left and 2k) and perpendicular (division axis perpendicular to fibre at mitosis, 18.9%, Fig. 2j right and 2k) divisions of myogenic cells on myofibres, in agreement with previous reports^35,38^. However, analysis of planar and perpendicular cell divisions during live imaging showed no significant differences in fate outcomes for WT (Fig. 2l) and *mdx* myogenic cells (Fig. S2g) that were related to the orientation of cell division plane.

Next, we quantified migration parameters of WT and *mdx* MuSCs by imaging *ex vivo*. While WT myogenic cells migrated actively, a fraction of *mdx* cells showed low migration capacity. We then stratified the data into ‘mobile’ and ‘static’ populations, to exclude static cells that can generate confounding effects when measuring speed and straightness. A migration speed threshold of 0.41 μm/min (Fig. S3l-m, see Methods) was used to discriminate between mobile and static fractions (Fig. S2h, Movie S4) for all *ex vivo* migration analyses (Fig. 2-4), and we observed that *mdx* myogenic cells exhibited a lower mobile fraction (mean = 0.19) than WT cells (mean = 0.52, p = 3.33e-19, Fig. 2m). Further, the mobile fraction of *mdx* myogenic cells showed a lower migration speed (mean: *mdx* = 36.4 μm/h, WT = 46.4 μm/h, p = 5.85e-12, Fig. 2n) and lower net displacement (Fig. S2i) than WT cells. The migration straightness was similar between the mobile fractions of WT and *mdx* cells (Fig. 2o), unlike our *in vivo* observations (Fig. 1h), albeit the *in vivo* analysis was performed on the total cell population. At the total population level (mobile + static), *mdx* myogenic cells showed a lower migration straightness *ex vivo* (mean: *mdx* = 0.15, WT = 0.21, p = 1.35e-03 Fig. S2j) and higher turning angle (mean: *mdx* = 97.6°, WT = 81.4°, p = 5.21e-13 Fig. S2k) than WT cells, as was observed *in vivo* (Fig. 1h,i).

Altogether, our *ex vivo* pipeline allowed high-throughput quantification and validation of precocious differentiation through SCD and impaired migration kinetics of *mdx* myogenic cells observed by intravital imaging during muscle regeneration.

### Cross-grafting assay uncouples cell-intrinsic fate and cell-extrinsic migration phenotypes in dystrophic MuSCs

Although DYSTROPHIN is a well-known structural protein of muscle fibres, it was reported to be expressed in myoblasts as a regulator of ACD^4^. Therefore, perturbed myogenic cell fate decisions and migration in *mdx* could result from lack of DYSTROPHIN in myoblasts and/or myofibres. To discriminate between intrinsic and extrinsic cues, we developed an *ex vivo* cross-transplantation assay where the behaviour of MuSCs of WT or *mdx* origin could be examined on myofibres of the opposite genotype and monitored continuously by live imaging. WT or *mdx* MuSCs from *Pax7^CreERT2^*; *R26^YFP^; Myog^ntdTom^* mice were engrafted to *mdx* or

WT receiving FDB myofibres (Fig. 3a-b and Fig. S5, hereafter referred to as ‘WT_to_*mdx* graft’ and ‘*mdx*_to_WT graft’). The transplantation of WT MuSCs to WT fibres (WT Ctrl) and of *mdx* MuSCs to *mdx* fibres (*mdx* Ctrl) were used to monitor possible effects of the grafting procedure (Fig. 2 and Fig. S2).

**Figure 3.**
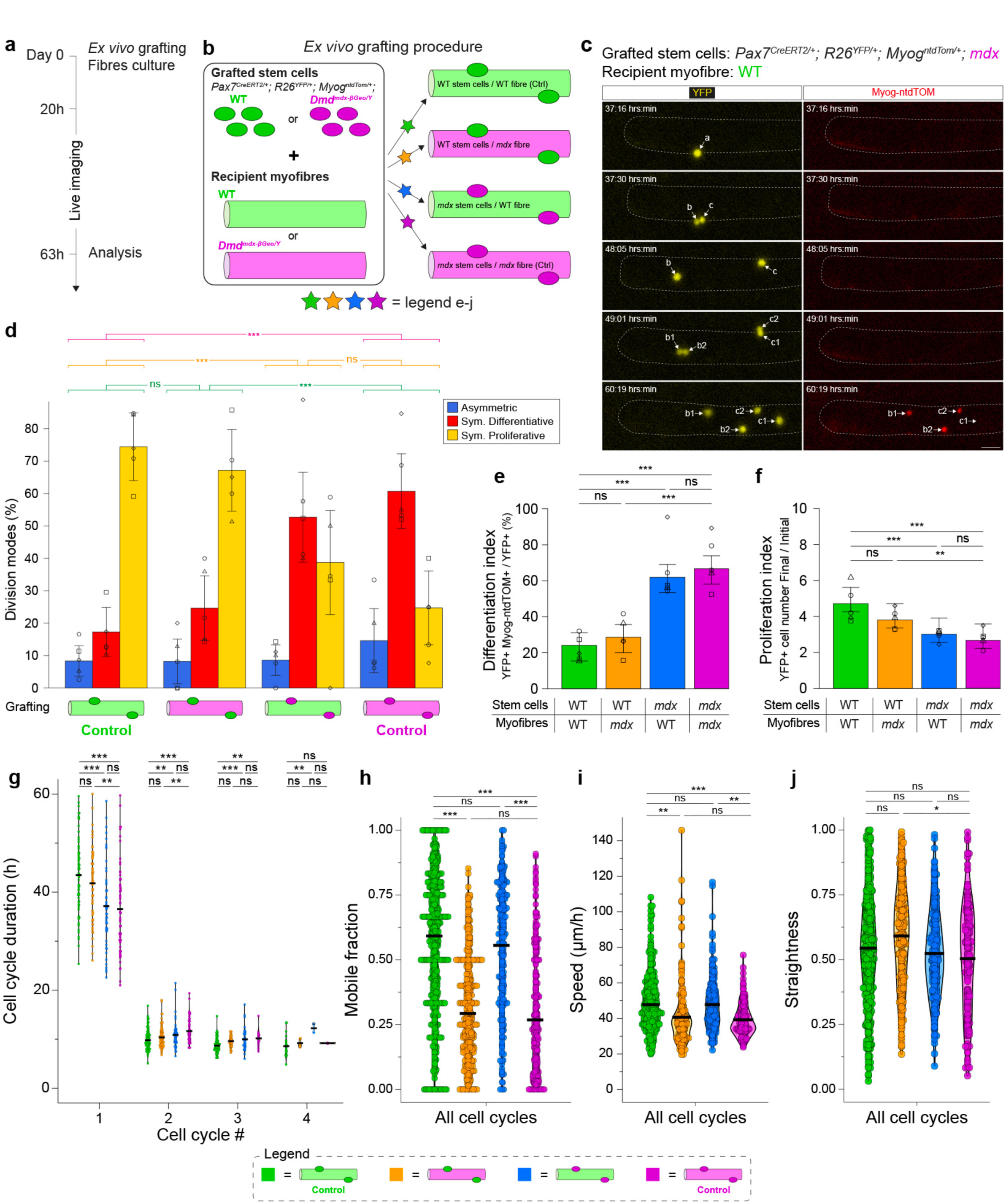
Cross-grafting assay uncouples cell-intrinsic fate and cell-extrinsic migration phenotypes in dystrophic MuSCs. **(a)** Experimental scheme. MuSCs and FDB fibres from 4-month-old WT and *mdx* male mice were isolated, cross-grafted (Fig. 3b), cultured and filmed in microwells for ∼ 63h. Also see Fig. S5. **(b)** Scheme of cross-grafting assay. MuSCs from *Pax7^CreERT2/+^; R26^YFP/+^; Myog^ntdTom/+^; Dmd^+/Y^*(WT) and *Dmd^mdx-βGeo/Y^* (*mdx)* 4-month-old male mice were isolated and grafted on FDB fibres from *Dmd^+/Y^* (WT) and *Dmd^mdx-βGeo/Y^* (*mdx)* 4-month-old male mice *ex vivo*, following 4 combinations: WT MuSCs on WT fibre (WT Ctrl), *mdx* MuSCs on *mdx* fibre (*mdx* Ctrl), WT MuSCs on *mdx* fibre (WT_to_*mdx*) and *mdx* MuSCs on WT fibre (*mdx*_to_WT). Also see Fig. S5. **(c)** Snapshots of a representative example of live imaging of a WT FDB fibre grafted with an *mdx* YFP-labelled MuSC. The grafted cell *a* divided to give rise to daughters *b* and *c*, which migrated and divided to give rise to granddaughters *b1*, *b2*, *c1* and *c2*. *Myogenin* expression was followed live with a Myog-ntdTOM reporter. Here, *b* executed a symmetric differentiative division, *c* executed an asymmetric division. Scale bar, 30 μm. See Movie S5. **(d)** Modes of cell divisions for all grafting conditions. N = 5 experiments. **(e)** Differentiation index for all grafting conditions. N = 5 experiments. **(f)** Proliferation index for all grafting conditions. N = 5 experiments. **(g)** Cell cycle duration (h) for all cell cycles for all grafting conditions. N = 5 experiments. Upon isolation and *ex vivo* culture, MuSCs exited from their niche and transited from below the basal lamina (sub-laminal) to above (supra-laminal) within 24h (Fig. S3a-c). MuSCs adhered directly to the myofibre basal lamina (supra-laminal position) after *ex vivo* grafting. The difference in cell cycle entry between WT and *mdx* MuSCs observed upon grafting (Fig. 3i) might reflect *mdx* MuSCs poised for activation, masked by a delay in niche exit in the endogenous condition (Fig. 2e). **(h)** Mobile fraction for all grafting conditions for merged cell cycles. N = 5 experiments. **(i)** Migration speed of mobile fraction for all grafting conditions for merged cell cycles. N = 5 experiments. **(j)** Migration straightness of mobile fraction for all grafting conditions for merged cell cycles. N = 5 experiments. Statistical tests: **(d)** Fisher test; **(e-f)** Wilcoxon test; **(g-j)** Linear mixed models. Horizontal lines in violin plot represent the mean. * *p* < 0.05, ** *p* < 0.01, *** *p* < 0.001.

Consistent with our observations above, myogenic cells divided mostly through proliferative symmetric divisions in the WT Ctrl graft (ACD = 8.3%; SCDd = 17.3%, SCDp = 74.4%, Fig. 3d), while they showed impaired SCDs in the *mdx* Ctrl control (ACD = 14.6%; SCDd = 60.7%, SCDp = 24.7%, p-value vs WT Ctrl = 1.09e-13), thereby validating the cross-transplantation assay (Fig. 2i). Comparison of WT Ctrl graft (Fig. 3d) with WT MuSCs on their own myofibre (Fig. 2i) showed a lower frequency of ACD (graft = 8.3%, endogenous = 12.9%) and SCDd (graft = 17.3%, endogenous = 32.5%) and higher frequency of SCDp (graft = 74.4%, endogenous = 54.2%), likely due to the lower sensitivity (short exposure times, slow maturation (t_0.5_ ∼ 1h) of tdTOMATO protein) of the Myog-ntdTOM live reporter compared to MYOG immunostaining). Notably, division modes in the WT_to_*mdx* graft (Fig. 3d) were like the WT Ctrl (p = 0.27) but different from the *mdx* Ctrl (p = 2.22e-09). Further, the division modes in the *mdx*_to_WT graft (Fig. 3d) were like the *mdx* Ctrl control (p = 0.12) but different from the WT Ctrl (p = 1.35e-08). These observations suggest that the division modes are driven primarily by MuSC-intrinsic cues and are not differentially affected by the myofibre niches.

Next, we measured differentiation and proliferation indexes (Fig. 3e-f) at the population level following cross-transplantations. As expected, the differentiation and proliferation indexes were different between the WT Ctrl and *mdx* Ctrl (p = 3.6e-11 and 5.8e-06 respectively). The differentiation index of the WT_to_*mdx* graft (Fig. 3e) was like the WT Ctrl (p = 0.62) but different from the *mdx* Ctrl control (p = 5.0e-09), and the differentiation index of the *mdx*_to_WT graft (Fig. 3e) was like the *mdx* Ctrl (p = 0.32) but different from the WT Ctrl (p = 7.5e-08). These results indicate that the precocious differentiation of *mdx* myogenic cells was not rescued by a WT myofibre, and that the differentiation of WT myogenic cells was not affected by the dystrophic environment. This was also the case for proliferation indexes which were found to be independent of the myofibre niche. Altogether, our results show that proliferation and differentiation decisions through ACD/SCDs are mostly driven by MuSC-intrinsic cues, and largely independent of the respective myofibre niches.

We then used *Myog^ntdTom^* live reporter mice to monitor cell fate decisions for each cell cycle over consecutive divisions (Fig. 3c and Movie S5). Most of the Myog-ntdTOM expression appeared ∼9h post-mitosis (Fig. S3e), with rare examples (10.2%) of Myog-ntdTOM-positive cells performing one more cell division (Fig. S3f), as reported^32^, and most SCDd cells originated from a Myog-ntdTOM-negative mother cell (89.8%; Fig. S3f).

We next measured cell cycle progression for all grafting conditions. Interestingly, *mdx* Ctrl cells executed their first division (mean 36.5h) before WT Ctrl cells (mean 43.5h, p = 1.32e-05, Fig. 3g), unlike WT and *mdx* MuSCs in their endogenous myofibre niche (Fig. 2e). *mdx* Ctrl cells cycled slower than WT Ctrl cells in subsequent cell cycles, as observed with the endogenous condition. Notably, WT_to_*mdx* cells had a cell cycle progression like WT Ctrl cells but distinct from *mdx* Ctrl cells, whereas *mdx*_to_WT cells had a cell cycle progression like *mdx* Ctrl cells but different from WT Ctrl cells (Fig. 3g), providing further evidence that proliferation kinetics of MuSCs is mostly driven by MuSC-intrinsic cues.

Finally, we measured migration parameters (mobile fraction, speed, straightness, displacement), for all or separated cell cycles (Fig. 3h-j and Fig. S3g-k). The mobile fraction (Fig. 3h), migration speed (Fig. 3i) and net displacement (Fig. S3j) of the WT Ctrl and *mdx* Ctrl differed (p = 4.69e-14, 9.68e-04 and 5.50e-07 respectively across all cell cycles), while their migration straightness (Fig. 3j and Fig. S3i) was similar when analysing the mobile fraction (p = 0.69 across all cell cycles) but different at the total population level (Fig. S3k, p = 2.48e-05 across all cell cycles), showing that cross-transplantations can recapitulate the differential migration phenotypes of WT and *mdx* MuSCs observed in their endogenous niches. Strikingly, the mobile fraction and migration speed of the WT_to_*mdx* graft were different from the WT Ctrl (p = 6.55e-14 and 4.05e-03 respectively across all cell cycles) but like the *mdx* Ctrl (p = 0.62 and 0.92 respectively across all cell cycles) (Fig. 3h-i). Conversely, the mobile fraction and migration speed of the *mdx*_to_WT graft were different from the *mdx* Ctrl (p = 3.24e-10 and 2.93e-03 respectively across all cell cycles) but like the WT Ctrl (p = 0.33 and 1.0 respectively across all cell cycles) (Fig. 3h-i). At the total population level, the migration straightness of the WT_to_*mdx* graft was like the WT Ctrl (p = 1.0) and the *mdx*_to_WT graft was like the *mdx* Ctrl (p = 0.64) (Fig. S3k). Altogether, these observations indicate that the migration properties of MuSCs, and their alterations in *mdx* mice, are largely determined by myofibre-derived signals.

### Excessive differentiation of *mdx* MuSCs is driven by the coordinated activity of p38 MAPK and PI3K pathways

To explore the molecular mechanisms behind the fibre-dependent migration defects in *mdx* MuSCs, we analysed RNA-seq data from FDB fibres of 2- and 5-month-old WT and *mdx* mice (Fig. 4a-c, Fig. S4a)^43^ and scRNA-seq data from gastrocnemius muscle of 2-month-old WT and *mdx* mice^7^ (Fig. S4b). Gene Ontology (GO) analysis revealed significant enrichment of terms related to migration and differentiation in *mdx* fibres and MuSCs (Fig. 4b, Fig. S4a-b). Differentially expressed genes included *Parva*, *Lrp1*, and *Adipoq* (negative regulators of migration^44–46^ and upregulated in *mdx*), and *Nexn* (positive regulator of migration^47^ and downregulated in *mdx*)(Fig. 4c). Additionally, *Tspan* and *Emilin1* (upregulated in *mdx*, Fig. 4c) and integrins (upregulated in *mdx*, Fig. 4b) were implicated in migration inhibition^48,49^.

**Figure 4.**
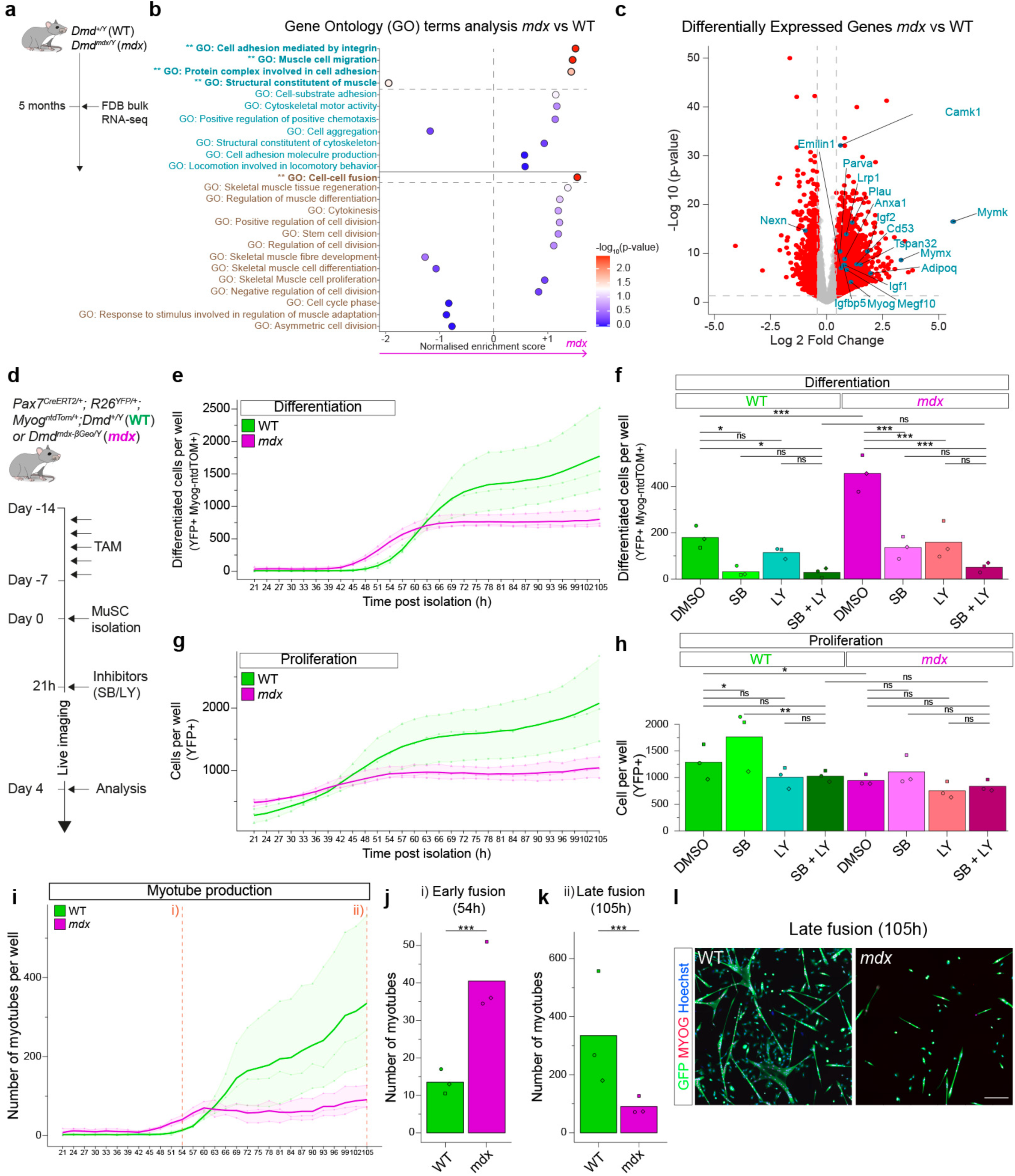
Excessive differentiation of *mdx* MuSCs is driven by the coordinated activity of p38 MAPK and PI3K pathways. **(a)** Pipeline of RNA-seq analysis of FDB fibres from 5-month-old *Dmd^+/Y^* (WT) and *Dmd^mdx/Y^* (*mdx*) mice^43^. **(b)** Gene Ontology (GO) terms enrichment analysis on RNA-seq analysis of FDB fibres from 5-month-old WT and *mdx* mice^43^. GO terms related to migration, proliferation and differentiation are shown. **(c)** Volcano plots of differentially expressed genes (DEGs) of FDB fibres from 5-month-old *mdx* vs WT mice. Genes highlighted in blue were selected from GO terms with significant enrichment in *mdx* vs WT fibres (Fig. 4b). **(d)** Experimental scheme. *Pax7^CreERT2/+^; R26^YFP/+^; Myog^ntdTom/+^; Dmd^+/Y^* (WT) and *Dmd^mdx-βGeo/Y^*(*mdx)* adult 4-month-old male mice were treated 5X with tamoxifen. YFP-labelled MuSCs from hindlimbs muscles were isolated by FACS, cultured and filmed for 4 days. Cells were treated with SB203580 (SB) and/or LY294002 (LY) from 21h after isolation. N = 3 experiments. See Movie S6. **(e)** Kinetics of differentiated cell production (number of YFP-positive Myog-ntdTOM-positive cells/well) of DMSO-treated WT and *mdx* cells. **(f)** Differentiated cells production of WT and *mdx* cells treated with SB and/or LY at 54h of culture. N = 3 experiments. **(g)** Kinetics of total cell production (number of YFP-positive cells/well) of DMSO-treated WT and *mdx* cells. **(h)** Total cell production of WT and *mdx* cells treated with SB and/or LY at 54h of culture. N = 3 experiments. **(i)** Kinetics of myotube production (number of myotubes/well) of DMSO-treated WT and *mdx* cells. Statistical comparisons were performed at 54h (j) and 105h (k). **(j)** Myotube production of DMSO-treated WT and *mdx* cells at 54h of culture. N = 3 experiments. **(k)** Myotube production of DMSO-treated WT and *mdx* cells at 105h of culture. N = 3 experiments. **(l)** Representative example of myotube production of DMSO-treated WT and *mdx* cells at 105h of culture. Scale bar = 200 μm. Statistical tests: **(f, h, j-k)** Linear mixed models. * *p* < 0.05, ** *p* < 0.01, *** *p* < 0.001.

To investigate the mechanisms of the precocious differentiation of *mdx* MuSCs, cells from hindlimb muscles of *Pax7^CreERT2/+^; R26^YFP/+^; Myog^ntdTom/+^;* WT and *mdx* mice were cultured with p38 MAPK and/or PI3K inhibitors (SB203580 [SB] and LY294002 [LY], respectively) and live-imaged continuously to track proliferation, differentiation, and fusion (Fig. 4d-l, Fig. S4c-e and Movie S6). As observed previously, vehicle-treated (DMSO) *mdx* cells showed earlier differentiation (Fig. 4e) and reduced proliferation compared to WT (Fig. 4g), leading to fewer cells (in total and differentiated).

In *mdx* cells, combined SB and LY treatment nearly abolished differentiation (88.9% reduction, p = 4.61e-07), while in WT cells, SB alone suppressed differentiation by 82.6% (p = 1.19e-02), with LY having no significant effect (Fig. 4f, Fig. S4c). These results indicate that *mdx* differentiation depends on both p38 and PI3K pathways, whereas WT differentiation relies primarily on p38.

SB-treated WT cells showed increased proliferation (p = 3.89e-02), while LY alone or combined with SB had no effect (Fig. 4h and Fig. S4d). In *mdx* cells, neither SB nor LY affected proliferation, indicating that the proliferation defect of *mdx* cells is not a direct consequence of their precocious differentiation but rather a distinct phenotype, as SB and/or LY treated *mdx* cells did not differentiate but still failed to proliferate.

Analysis of myotube production over time revealed that *mdx* cells generated myotubes earlier but failed to sustain production at later stages (Fig. 4i-l and Fig. S4e). Blocking p38 or PI3K pathways did not rescue this defect (Fig. S4e). *Ex vivo* fibre experiments confirmed the role of the p38 pathway, as SB treatment rescued *mdx* differentiation to WT levels without affecting motility (Fig. S4f-j).

In summary, *mdx* MuSCs exhibit p38- and PI3K-dependent precocious differentiation, contrasting with WT cells’ p38-only dependence, together with proliferation defects. These intrinsic defects lead to premature myotube formation and insufficient production of fusion-competent cells at later stages.

## Discussion

Using unique *in vivo* and *ex vivo* pipelines, we show that dystrophic MuSCs exhibit impaired migration kinetics driven by the dystrophic myofibre niche, as well as proliferation defects and precocious differentiation through symmetric divisions. These processes are largely cell-autonomous, rely on p38 and PI3K signalling, and result in inefficient myotube production. Notably, we demonstrate that fate decisions are largely uncoupled from cell motility.

The regulation of dystrophic MuSC fate and its contribution to impaired regeneration in DMD has been long debated, with conflicting studies reporting decreased or increased differentiation, proliferation, and fusion defects^4,9,50,51^. We propose that these discrepancies arise in part from static and discontinuous temporal analyses, as well as varying models. By integrating dynamic *in vivo* imaging with *ex vivo* assays, we resolve inconsistencies and highlight the importance of assessing cell fate decisions continuously and temporally, particularly in the case of DMD. We demonstrate premature differentiation and proliferation defects in *mdx* MuSCs, which fail to sustain myotube production at later stages. These findings align with the concept of secondary MuSC-opathies in DMD^2^.

We further identify PI3K and p38 as key drivers of precocious differentiation in *mdx* MuSCs, consistent with their convergence for terminal differentiation^52^. As insulin-like growth factor I (IGF-I)-PI3K-Akt and p38 signalling are influenced by inflammation, abundant in dystrophic tissues, immune cells likely contribute to this dysregulation^53,54^. Notably, p38 inhibitors like Losmapimod (GW856553, p38α/β inhibitor) have shown promise in clinical trial for Facioscapulohumeral muscular dystrophy (FSHD) through DUX4 inhibition and for COVID-19 patients^55^, while PI3K/Akt inhibition reportedly expands PAX7+ MuSCs in DMD^17^, highlighting their therapeutic potential.

While asymmetric cell division has been well studied in the myogenic lineage^3,4,35^, the role of symmetric divisions remains unexplored although they have major impact on cell population dynamics. We show here that the dynamics of transit-amplifying myogenic cells are dominated by symmetric divisions (∼ 10-15% ACD), with unbalanced symmetric proliferative and differentiative divisions contributing to the dystrophic phenotype. Although prior work suggested impaired ACD in *mdx* MuSCs^4^, our findings highlight defects in symmetric divisions, potentially reflecting differences between early activation and transit-amplifying stages or static versus dynamic analyses.

Motility defects of dystrophic myoblasts have been noted *in vitro*, linked to reduction in migration and aberrant matrix interactions^18,56^. Further, aberrant migration of myoblasts outside regenerating fibres has been proposed to result in branched myofibres^23^, a sign of failed regeneration in dystrophic muscles^29^. Our study provides the first *in vivo* evidence of impaired migration, driven largely by the dystrophic niche. Delayed separation after mitosis and immobile myogenic cells likely disrupt proper cell distribution along regenerating fibres, contributing to failed and heterogeneous repair^57^. Enhancing migration through small molecules like Sdf1, HGF, and FGF, or by modulating the niche^18^, represents a promising avenue for improving regeneration in DMD.

In conclusion, our findings reinforce the notion that DMD is not solely a myofiber disease but also a MuSC disease, driven by complex and dynamic defects. This has significant implications for regenerative medicine, particularly gene therapies targeting MuSCs to restore long-term stem cell function and tissue homeostasis.

## Methods

### Mouse strains and genotyping

*Pax7^CreERT2^*^26^, *R26^mTmG^*^27^, *R26^YFP^*^33^, *Myog^ntdTom^*^32^ and *Dmd^mdx-βGeo^*^28^ mouse breeding was performed on a mixed B6D2F1/JRj background. 4-month-old male littermates were used, hemizygous for *Dmd* and heterozygous for other alleles. Animals were handled according to national and European community guidelines, and protocols were approved by the ethics committee at Institut Pasteur (Licence 2015-0008 and DAP 220077). Mice were genotyped at 3 weeks by PCR from an ear punch biopsy. PCR primers and conditions are listed in Table S1.

### Tamoxifen treatment and muscle injury

Tamoxifen (T5648, Sigma) was reconstituted at 25 mg/ml in corn oil/ethanol (2.5%) and stored at −20°C. Tamoxifen was administered (200 μl/injection) by intragastric injection, once or 5X over 5 days, followed by 1 week of chase.

Muscle injury was done as described^58^. Briefly, mice were anesthetized with ketamine (Imalgene 1000^®^, 100 mg/kg in NaCl 0.9%) and xylazine (Rompun 2%^®^, 12 mg/kg in NaCl 0.9%). The FDB muscle was injected with 20 μl of 10 μM cardiotoxin (L8102, Latoxan) in NaCl 0.9% using a 30G insulin syringe (BD, Micro-Fine 324826).

### Intravital imaging

4-month-old male littermates *Pax7^CreERT2/+^; R26^mTmG/+^; Dmd^+/Y^*or *Dmd^mdx-βGeo/Y^* were treated once with tamoxifen, followed by 1 week of chase, and FDB muscles were injured (see below). FDBs of WT and *mdx* mice were injured one day apart as intravital imaging sessions allowed to process 1 mouse/day.

On the day of imaging, the mouse was treated with buprenorphine (intraperitoneal injection, 0.1 mg/kg, Vetergesic^®^). After 30 min, the mouse was anaesthetised in an induction chamber with isofluorane (Iso-vet) 3% and O_2_ 2 L/min in an incubation chamber, transferred onto a heat pad (Minerve) and anaesthesia was maintained using a rodent-adapted mask (Minerve) with 1-1.5% isoflurane and O_2_ 0.6 L/min delivered through a gas humidifier. Ophthalmic gel (Ocry-gel^®^) was applied to the eyes to prevent corneal drying. The mouse foot was stabilised with tape, cleaned with 70% ethanol, and a 2 x 3 mm area of skin was carefully removed to expose the FDB. The FDB was immediately gently pressed against a glass-coverslip attached (silicone) to a 3D-printed custom coverslip holder, immobilised with tape and sealed (silicone) to the coverslip. After polymerisation of the silicone, the foot was turned upside-down to have the coverslip and the FDB facing up, and the coverslip holder was secured to the heating pad (silicone). The mouse was rehydrated with pre-warmed 0.9% NaCl (0.5 ml per hour) through a subcutaneous catheter (SV*S23NL30, Terumo) (Fig. S1). Imaging was done using a multiphoton microscope (TrimScope Matrix, Miltenyi Biotec). Excitation was performed at 950 nm with 3% laser power (Insight X3, Spectra Physics) and photons were collected with a 25x/NA 1.05 water immersion objective (Evident, Olympus) and detected on Gaasp detectors (Hammamatsu) at 510/20 nm (GFP) and 580/25 nm (tdTOMATO).

Imaging was performed by capturing tiles of 393 x 393 μm (1024 x 1024 pixels) on a 2.2 mm x 0.8 mm mosaic, with a z-stack of 80 μm (z-step 3 μm). Images were acquired every 10 min for 8-10h. Mouse breathing was monitored every hour and anaesthesia adapted if necessary. At the end of the imaging session, the mouse was euthanised by cervical dislocation. Assembling of movies was done with ImageJ with custom-made macros and the Stitching plugin^59^.

### MuSC analysis, isolation, culture and live-imaging

FDB or hindlimb muscles were dissected and minced in a drop of cold collagenase type 2 (1000U/ml, WOLS04177, Serlabo) in Muscle Dissociation Buffer (MuDB, Nutrient mixture Ham F10 (N6635, Sigma), 10% heat-inactivated Horse serum (HS, 11510516, ThermoFisher), supplemented with NaHCO_3_ to pH7.4 in distilled water). Samples were incubated in 10 ml of a collagenase 2/MuDB for 90 min at 37°C in a water bath under gentle agitation (70 rpm). Samples were mechanically dissociated (25 ml pipette), MuDB was added up to 50 ml and samples were spun (10 min, 500 g at RT). The supernatant was removed up to 20 ml, cell pellet was resuspended and 1 ml of Dispase (1.8 U/ml in MuDB, 17105-041, Gibco) and 20 μl DNase I (10 mg/ml in DMEM, 11284932001, Roche) were added. Samples were incubated for 90 min at 37°C in a water bath under gentle agitation (70 rpm). Samples were passed through a syringe (Agani Needle 20G, 050109B Fisher Scientific) 10 times, filtered through 40 μm strainer (352235, Dutscher), spun (10 min, 500 g at 4°C) and resuspended in 500 μl of DMEM/2% HS before cytometry.

Using Fluorescence Activated Cell Sorting (FACS), samples were analysed (CytoFLEX, Beckman Coulter) and MuSCs were isolated (Aria III, BD Biosciences) based on cell size, granularity and YFP fluorescence (70 μm nozzle). Cells from *Pax7^CreERT2/+^; R26^YFP/+^* mice were used to determine the positivity threshold of Myog-ntdTOM. Cells were collected in differentiation medium (88.5% DMEM, 10% HS, 1% Penicillin/Streptomycin, 0.5% Chick Embryo Extract (MD-004D-UK, Life Science Production)) at 4°C.

Isolated MuSCs were plated at 4500 cells/cm^2^ in differentiation medium in 96-well plates (PhenoPlate (6055300, Revvity), well surface = 0.32 cm^2^) coated with fibronectin (F1141, Sigma; 40 μg/ml in NaHCO_3_ 0.1 M pH8.3 for 30 min at RT followed by 3 PBS washes). p38 MAPK inhibitor (SB203580 5 μM, 559389, Sigma), PI3K inhibitor (LY294002, 10 μM, 9901S, Cell Signaling) or vehicle (DMSO) were added at 21h post plating. Cells were incubated at 37°C, 3% O_2_, 5% CO_2_.

Cells were live-imaged with a Zeiss Observer Z1 equipped with a Plan-Apochromat 20x/0.8 M27 objective, Colibri 7 LEDs, a Hamamatsu Orca Flash 4 camera and piloted with Zen software (Carl Zeiss), at 37°C, 5% CO_2_ and 3% O_2_ (Pecon incubation chamber). Each well was imaged every 3h in 2 channels (YFP, DsRed).

### Microwell plate assembly

The microwell plate is composed of two Plexiglas plates (bottom plate: 128 mm x 85 mm x 10 mm; top plate: 123 mm x 81 mm x 10 mm) in which 4 rectangles (55.5 mm x 34 mm x 10 mm) have been laser-cut (Speedy 300, Trotec), cleaned and attached together (Fig. S2a) with silicone (MoldStar 20T, Smooth-on). 4 glass coverslips (Claritx Coverglass #1, Eloïse-SARL) were cut to 58 mm x 38 mm, cleaned and sealed (Fig. S2a) with silicone to the outer part of the bottom Plexiglas plate to form 4 glass-bottom chambers. The lid of a 6-well dish (TPP) was used to close the microwell plate on the top Plexiglas plate.

Silicone sheets with microwells were cleaned and placed on glass coverslips (Fig. S2a) and fixed to the glass with additional silicone on the edges. Microwells were pierced manually (biopsy ear punch in 1 mm-thick silicone; microwell volume ∼ 3 μl. 50 microwells/glass-bottom chamber. Fig. 2b and Fig. S2a) or formed using a 3D-printed mold (3 mm-thick silicone; microwell volume ∼ 6 μl. 200 microwells/glass-bottom chamber Fig. S2a).

Spinning-disk confocal live-imaging (Fig. 2c and Movie S3) was carried out using a 2 well glass bottom (80287, Ibidi) in which two silicone sheets with microwells were sealed with silicone.

All the material (Plexiglas plates, coverslips, silicone microwells) was cleaned and sterilized with soap, water and ethanol 70% before use. After assembly of the microwell plate, microwells were washed with PBS and equilibrated with culture medium (5 ml per chamber, final volume 10 ml) to prevent adhesion of muscle fibres. Air bubbles in microwells were removed with a P200 pipette and single FDB fibres were loaded in microwells. As the height of the microwells (1 or 3 mm) is 20-60 times larger than the thickness of an FDB fibre (50 μm), this microwell design allows confinement, culture and live-imaging of single FDB fibres in non-adherent conditions.

All subsequent manipulations after fibre loading (medium exchange, live-imaging, immunostaining) were done as usual when manipulating a 6-well plate.

### Single myofibre isolation, culture and live-imaging

A solution of 0.2% collagenase Type 1 (C0130, Sigma) was prepared in DMEM GlutaMAX (31966, ThermoFisher), filtered (0.22 μm) and kept at 37°C. Mice were sacrificed by cervical dislocation. The foot was cleaned with 70% ethanol and the skin covering the *Flexor Digitorum Brevis* (FDB) was cut from the base of the heel to the toes. The FDB muscle was dissected away from the surrounding tissue, being handled carefully by the proximal tendon to avoid muscle damage and contraction. The FDB was incubated in 5 ml of collagenase solution in Sterilin 7mL Polystyrene Bijou Containers (11399133, Fisher) for 2.5 h at 37°C.

Individual muscle fibres were released by mechanical dissociation (glass Pasteur pipette) of FDB in Sterilin petri deep dishes (10655821, Fisher) filled with pre-warmed DMEM. The majority of the FDB fibres were released after five rounds of 10 vigorous but careful back-and-forth pipetting, separated by 5-min pauses at 37°C. After each round, the FDB was transferred to a new Sterilin dish with pre-warmed DMEM for further dissociation. Large debris were removed with a plastic Pasteur pipette. When most fibres were dissociated, two-thirds of the DMEM volume were removed from each Sterilin dish. Muscle fibres were then pooled in a new Sterilin dish filled with pre-warmed DMEM, washed with DMEM and distributed to 5 Sterilin dishes containing culture medium and caps of 15 ml Falcon tubes in their centre (fixed with silicone) to prevent fibres from concentrating and aggregating in the centre of the dishes. Sterilin dishes and Pasteur pipettes were coated with horse serum (HS, 11510516, Fisher Scientific) before use to prevent adhesion of muscle fibres. Muscle fibres were loaded in the microwell plate in bulk and/or manually under the binocular loupe using a P20 pipette.

Muscle fibres were cultured in differentiation medium at 37°C, 5% CO_2_, 3% O_2_. For p38 inhibition experiments, p38 MAPK inhibitor (SB203580 5 μM, 559389, Sigma) or vehicle (DMSO) were added directly to the culture medium ∼39 h post-isolation of muscle fibres.

Spinning-disk confocal live-imaging of FDB fibres (Fig. 2c and Movie S3) was performed immediately after isolation using a Spinning Disk Confocal Yokogawa CSU-W1 on a Nikon Ti2E equipped with a 20x dry objective (numerical aperture 0.75, working distance 1 mm), a motorized XY stage (with a Z piezo stage, 200 μm range), a Hamamatsu Orca Flash 4 camera (pixel size 6.5 μm, 2048 × 2044 pixels, quantum efficiency 82%), and piloted with Nikon software (NIS Element) at 37°C, 5% CO_2_ and 3% O_2_ with humidity control and objective heater (Okolab). Each fibre was imaged every 10 min in 488 nm (300 ms, 1.5% laser power) and 561 nm (200 ms, 0.8% laser power), with a Z-stack of 71 μm (1 μm slices).

Widefield live-imaging of FDB fibres (Fig. 2d-o, Fig. S2, Fig. 3 and Fig. S3e-m) in microwell plates was performed between ∼17 h and 64 h post-isolation with a Zeiss Observer Z1 (see above). Each fibre was imaged every 10-14 min in 2 or 3 channels (Brightfield, YFP, DsRed), with a Z-stack of 50 μm (5 μm slices). Stage parameters were set at 50% speed, 5% acceleration.

### Immunocytochemistry

Muscle fibres were fixed with 4% paraformaldehyde (PFA, 15710, Euromedex) in PBS (D1408, Sigma) for 10 min at room temperature (RT), washed twice 5 min in PBS, permeabilized in cold permeabilization buffer (Hepes 20 mM pH 7.4, MgCl_2_ 3 mM, NaCl 50 mM, Sucrose 300 mM, 0.5% Triton X-100) for 15 min at 4°C and washed three times in PBS. Fibres were blocked with 10% Goat Serum (GS, 11540526, ThermoFisher) in PBS (0.22 μm-filtered) for 1h at RT and washed once in 2% GS in PBS (0.22 μm-filtered). Excess of liquid between microwells was removed, leaving each microwell with 3 μl of 2% Goat Serum solution. Primary antibodies (Table S2) were diluted in 2% GS to a 4x concentration compared with their final concentration. 1 μl of diluted antibodies was added to each microwell (3 μl) and incubated overnight at 4°C. After three washes with 2% GS for 5 min at RT, the excess of liquid was removed. Secondary antibodies (Table S2) were diluted in 2% GS to a 4x concentration compared with their final concentration. 1 μl of diluted antibodies was added to each microwell (3 μl), for 1h at RT. Fibres were washed with PBS (5 min at RT), incubated with 1 μg/ml Hoechst 33342 (H1399, ThermoFisher) in PBS (5 min at RT), washed twice with PBS (5 min at RT) and stored at 4°C.

Immunostainings were analysed with Zeiss LSM800 confocal microscope. Each fibre was imaged in 3 to 5 channels (Brightfield, 488, 555, 633, Hoechst), with a centred Z-stack of 42 μm (1.9 μm slices) with a Plan-Apochromat 20X/0.8 M27 objective.

Our confinement system allowed filming and staining the same fibres (Fig. 2a-b, Fig. S2a and Movie S3), despite their culture in floating conditions.

### Cross-transplantation assays

Two 4-month-old male littermates *Pax7^CreERT2/+^; R26^YFP/+^; Myog^ntdTom/+^; Dmd^+/Y^* or *Pax7^CreERT2/+^; R26^YFP/+^; Myog^ntdTom/+^; Dmd^mdx-βGeo/Y^* were treated 5X with tamoxifen, followed by 1 week of chase, and served as donors of WT or *mdx* YFP-labelled MuSCs. The *Myog^ntdTom^* allele harbours a nuclear tdTOMATO reporter for *Myogenin* expression, in the 3’UTR of the *Myogenin* locus. Two 4-month-old male littermates *Dmd^+/Y^* or *Dmd^mdx-βGeo/Y^* served to generate WT or *mdx* recipient FDB fibres (Fig. 3b and S5).

For each recipient mouse, fibres were isolated (see above) from 1 FDB. Half of the isolated recipient fibres were split into two Sterilin dishes (with caps of 15 ml Falcon tubes fixed in the centre and culture medium) and then transferred into two 15 ml Falcon tubes with the base cut (to form a cylinder) and attached to the lid of a Sterilin dish (silicone). This setup allowed to load a large volume (∼ 12 ml) in the Falcon tube and to concentrate the recipient fibres in a small surface (cross section area of 15 ml tube, ∼ 1.8 cm^2^) after sedimentation to maximise contact between recipient fibres and grafted MuSCs (see below). The recipient fibres were then incubated at 37°C, 5% CO_2_ and 3% O_2_ while the following steps were carried out.

For each donor mouse, fibres were isolated (see above) from both FDBs. Fibres were washed twice in PBS, PBS was removed as much as possible and TrypLE Express (12604013, ThermoFisher) was added (2 ml) for 15 min at 37°C. MuSCs were then dissociated mechanically using a P1000 pipette, filtered through a 30 μm filter (130-041-407, Miltenyi) into a 15 mL falcon, centrifuged at 500g for 15 minutes at 21°C and resuspended in 500 μl differentiation medium.

The medium in the 15 ml Falcon tubes containing the recipient fibres was removed to leave 1.5 ml, to which 250 μl of MuSCs from donor mice were added. WT and *mdx* MuSCs were incubated with both WT and *mdx* fibres, corresponding to 4 grafting conditions. MuSCs and recipient fibres were incubated for 5 h at 37°C, 5% CO_2_ and 3% O_2_. For each grafting condition, medium was added (10 ml) in the 15 ml Falcon tube, the cell suspension was transferred into two Sterilin petri dishes to facilitate handling and the fibres were loaded in microwells of 1 chamber of a custom-made microplate (see above). Fibres were loaded in bulk from one Sterilin dish and manually from the other to reach full occupancy of microwells. During loading of a chamber, the other chambers were covered with Parafilm to ensure sterile conditions.

The microplate was then incubated at 37°C, 5% CO_2_ and 3% O_2_ before live-imaging. Under these conditions, each chamber (*i.e.* grafting condition) of the microwell plate contained ∼ 200 recipient fibres isolated in microwells, ∼ 25% of them grafted with 1-2 YFP-labelled MuSCs from donor mice. For each experiment, ∼ 20 fibres per grafting condition were imaged live.

### *Ex vivo* cell tracking and modes of cell division

Widefield live-imaging data of MuSCs on FDB fibres *ex vivo* was converted to 8-bit Tiff with ImageJ (https://github.com/gletort/ImageJFiles/blob/master/convertFiles/convertCZIto8bTiff.ijm). MuSCs were tracked manually using Phase and YFP channels for all timepoints with Trackmate^60^, and data was exported to .xml format for further analyses.

The fate of sister MuSCs, identified by cell tracking, was determined by immunostaining at the end of the experiment (Fig. 2f-g,i,l and Fig. S2g) or continuously using live-imaging and the Myog-ntdTOM reporter (Fig. 3d-f and Fig. S3f).

Divisions were classified as symmetric proliferative (both sister cells MYOG or Myog-ntdTOM-negative), symmetric differentiative (both sister cells MYOG or Myog-ntdTOM-positive) or asymmetric (one sister cell MYOG or Myog-ntdTOM-positive, the other MYOG or Myog-ntdTOM-negative).

The orientation of cell division was classified according to the position of sister cells at mitosis with respect to the FDB fibre: parallel if both sisters were in contact with the fibre, perpendicular if one of the sister cells had no contact with the fibre at mitosis.

### Analysis of cell migration

The migration of MuSCs was analysed *in vivo* (Fig. 1f-l) and *ex vivo* on muscle fibres (Fig. 2m-o, Fig. S2h-k, Fig. 3h-j and Fig. S3g-j). We first corrected the movements of regenerating muscles or FDB fibres to analyse MuSC migration in a static referential.

Displacement (translation, rotation, deformation) of FBD fibres *ex vivo* (due to culture in floating conditions) was corrected using a newly developed Fiji plugin: cellsOnFiber (distributed under the BSD-3 license and available in open source: https://gitlab.pasteur.fr/gletort/cellsonfiber/). Inputs are 3D movies of FDB fibres with TrackMate tracks of their associated MuSCs at every timepoints. This plugin detects the fibre from 2D projection of transmitted light images at each time frame with a specialized neural network, aligns it on the segmented fibre from the previous frame and corrects the TrackMate tracks accordingly. Outputs are 3D movies of FDB fibres and MuSCs TrackMate tracks corrected for fibre displacement.

*Training dataset*: We first imaged FDB fibres from *R26^mTmG^* mice in both the fluorescent channel (DsRed) and transmitted light and we obtained a ground-truth of fibre segmentation by applying a simple threshold on the fluorescent channel. The dataset was composed of 97 filmed scenes imaged on 149 time points. For the training, we subsampled the dataset every 20 timepoints for each scene. Overall, around 750 images of size 352 x 352 pixels of fibre in transmitted light with its ground-truth based on the thresholded fluorescent channel were used for training with basic data augmentation (flippings, translation and histogram). The dataset was randomly split so that 20% of the fibres were used for validation and the rest for training. Test was performed visually on dataset without ground-truth.

*Network architecture and training:* We used a U-Net architecture^61^ to detect the fibre contour from transmitted light 2D images. After testing different size of the network, we built a U-Net with 5-layer blocks in the left part (condensing part) of the network and 16 initial features in the first layer. The implementation was done with the python Keras library. The loss was calculated with the Jaccard distance, and the final score used was the intersection over union (IOU). Training was performed on a local computer with 1 GPU, for 30 epochs with a batch size of 30. The trained network is accessible here: https://gitlab.pasteur.fr/gletort/cellsonfiber/-/blob/main/networks/FiberFromTransMatch.zip.

*Registration and track correction:* The plugin calculates the registration to apply to the movie on the segmented fibre images to obtain a stabilised fibre. The fibre was segmented in each Z-slice of the time frame and projected in 2D. The transformation to register the fibre is calculated in our plugin with TurboReg plugin^62^ on the projected movie of the segmented fibre. The resulting transformation as then applied to each Z-slice of the movie of each input channel and used to correct the TrackMate trajectories to the new registered movie. The obtained tracks correspond to the cell motion relative to the immobilised fibre in 3D.

*Limitations:* In 7% of the movies, FDB fibre movement or deformation impaired analysis, and the plugin could not correct for this motion. These tracks were not used in any motility analysis but could still be used in lineage analysis (e.g. cycle duration, modes of divisions).

To correct for movements of the tissue during intravital microscopy, a first step of image registration was performed using ∼ 50 fixed points (mostly blood vessels) manually tracked with TrackMate Manual tracking^60^. We calculated the image registration on the channel containing the tissue information and applied it to channel containing the MuSCs. 3D registration was calculated with itk-elastix python module^63^. For each time frame, the registration was iteratively calculated to align to the previous frame. The registration was optimised in this step to decrease the distance between the same point at two consecutive times. The rigid registration was applied at 4 levels of resolution with 1000 iterations and a size of the final spacing grid of 50 pixels. This first registration was used to compensate for large local translation/rotation. Then a finer step was applied with b-spline based transformation to account for local deformation of the tissue, again with itk-elastix library. This step did not use the reference point and calculated the registration based on the intensity Mutual Information. The spline registration was performed at 2 resolution levels, with 1000 iterations and a size of the final spacing grid of 200 pixels. Finally, the calculated transformations were applied on all the channels of the original movie. All these steps were implemented in a Napari plugin napari-3dtimereg (distributed under the BSD-3 license and available in open source here: https://gitlab.pasteur.fr/gletort/napari-3dtimereg).

Next, we aimed at analysing the speed and directionality of migrating cells. Cell trajectories and lineage information were extracted from TrackMate files with a customed Jython script. The trajectories were then analysed with R scripts to measure different motility parameters for each cell.

Some MuSCs did not actively migrating on *ex vivo* FDB fibres, notably for *mdx* MuSCs, and some cells exhibited irregular trajectories, with phases of very small movements around the same point and more effective phases with more directional and faster motion. To analyse migration parameters on actively migrating cells, we fractionated the population into mobile and static subsets (Fig. S3l-m). The threshold value between mobile and static was determined by analysing the turning angle (see below) and the instantaneous speed (see below) between each consecutive time frames for WT and *mdx* MuSCs in the endogenous (dataset of Fig. 2 and S2) (Fig. S3l) and grafting condition (dataset of Fig. 3 and S3, control conditions WT Ctrl and *mdx* Ctrl) (Fig. S3m). WT cells (pooled from endogenous and WT Ctrl grafting datasets) showed two subpopulations (mobile with high speed/low turning angle and static with low speed/high turning angle), with a median log speed of 0.41 μm/min allowing to discriminate these subpopulations. *mdx* cells (pooled from endogenous and *mdx* Ctrl grafting datasets) showed mostly one population (low speed/high turning angle), with a median log speed of 0.18 μm/min. We then used a threshold value of 0.41 μm/min to discriminate between mobile and static subsets for all *ex vivo* datasets (Fig. S3l-m). Intravital imaging during muscle regeneration also identified some cells with low migration capacity, notably for *mdx* mice. Nevertheless, we could not identify a specific parameter to discriminate clearly mobile and static subsets. The analysis of cell migration properties *in vivo* was then performed on the total population of cells.

Turning angle (°) measures the average angle between two consecutives cell displacements (Fig. 1f). This measure varies from 0° (straight in the forward direction) to 180° (backward direction). The average speed (μm/h) is the ratio between the total distance travelled and the trajectory duration. The net distance (μm) measures the effective distance (3D) between the first and the last point of the trajectory. The total distance (μm) measures the total distance (3D) travelled by the cell as the sum of the distances between each time frames. The straightness is the ratio of the net distance over the total distance (Fig. 1f). If the trajectory was totally straight, then these two parameters are equal and the straightness is equal to 1. Otherwise, the cell had travelled more in total (TotalDistance) than the observed resulting distance (NetDistance), so the straightness is closer to 0.

For *in vitro* experiments, motility parameters were analysed for separated and pooled cell cycles.

### *In vitro* analysis of proliferation, differentiation and myotube production

Raw ZEN .czi files were analysed with ImageJ using the threshold tools to segment cells at each timepoint. Proliferation was assessed by measuring the number of YFP-positive objects per timepoint, differentiation by measuring the number of YFP and Myog-ntdTOM-positive objects, and myotube production by measuring the number of YFP-positive cells with an aspect ratio (Max diameter/Min diameter) superior to 2.

### Statistical analyses

For cell migration, sister cell separation *in vivo* and *in vitro* analysis, linear mixed models were fitted using the nlme R package and including the microwell identifier (nested within the experiment number) as random effects to test for the genotype [resp. inhibitor] factor. The mixed effects models included specific cell cycle variances when appropriate. The cell cycle variable was included in interaction with the genotype [resp. inhibitor] when testing for this effect for each cell cycle separately, otherwise the cell cycle was included as an additive main fixed effect. Pairwise comparisons were extracted from the linear mixed models using the emmeans R package and adjusted for multiple testing using the Tukey’s method.

For FACS and *ex vivo* analysis, mean comparisons were performed using unpaired two-tailed Wilcoxon tests unless stated otherwise. Comparisons between more than two groups were performed using Kruskall-Wallis tests followed by a post-hoc pairwise comparison.

### RNA-sequencing analyses

For analysis of bulk FDB fibres RNA-seq^43^, raw fastq files were downloaded from the GEO repository GSE162455 and aligned to the mm10 reference genome using STAR (v2.7.9a)^64^. Mapped reads were then quantitated using the RSEM pipeline (v1.3.3)^65^. Quality control plots and differential expression analyses were performed with the R package DESeq2 (v1.36.0)^66^.

For analysis of single-cell RNAseq^7^, Seurat objects were obtained from the authors and differentially expressed genes between conditions in MuSCs were identified using Wilcoxon tests implemented in the FindMarkers function from the R package Seurat (v4.4.0)^67^.

Gene Set Enrichment Analyses were performed with R using the fgsea function with default parameters from the *fgsea* package (v1.24.0). For each Gene Ontology term included in the analysis, genes included in the GO term and its children’s terms were assigned to the specific term. GO terms with less than 10 genes were discarded. Genes were ranked using −log10(adjusted p-value) * log2(fold change).

## Abbreviations

2D: two-dimensional
3D: three-dimensional
ACD: Asymmetric Cell Division
Ctrl: Control
CO2: Carbon dioxide
DEG: Differentially Expressed Gene
DMD: Duchenne Muscular Dystrophy
DMEM: Dulbecco’s Modified Eagle medium
DMSO: Dimethyl sulfoxide
dpi: days post-injury
DsRed: Discosoma Red Fluorescent Protein
FACS: Fluorescence Activated Cell Sorting
FDB: Flexor Digitorum Brevis
GEO: Gene Expression Omnibus
GO: Gene Ontology
GRMD: Golden Retriever Muscular Dystrophy
GS: Goat Serum
HS: Horse Serum
IGF-I: Insulin-like Growth Factor I
IOU: Intersection Over Union
MAPK: Mitogen-Activated Protein Kinases
mGFP: membrane-Green Fluorescent Protein
MuSC: Muscle Stem Cell
MYF5: Myogenic Factor 5
MYOD: Myoblast Determination Protein 1
MYOG: Myogenin
Myog-ntdTOM: Myogenin nuclear tdTOMATO live reporter
NaCl: Sodium chloride
O2: Oxygene
PAX7: Paired box 7
PBS: Phosphate Buffered Saline
PCR: Polymerase Chain Reaction
PFA: Paraformaldehyde
PI3K: Phosphoinositide 3-kinases
PTEN: Phosphatase and tensin homolog
RNA-seq: Ribonucleic Acid Sequencing
RT: Room Temperature
SCD: Symmetric Cell Division
SCDd: Symmetric Cell Division differentiative
SCDp: Symmetric Cell Division proliferative
scRNA-seq: single cell Ribonucleic Acid Sequencing
TAM: Tamoxifen
TOM: Tomato
WT: Wild Type
YFP: Yellow Fluorescent Protein

## Acknowledgements

We acknowledge funding support from the Institut Pasteur, Agence Nationale de la Recherche (Laboratoire d’Excellence Revive, Investissement d’Avenir; ANR-10-LABX-73 to ST), Association Française contre les Myopathies (#23774 to BE), European Research Council (Advanced Research Grant #101055234 to ST), La Fondation ARC pour la Recherche sur le Cancer and the Centre National de la Recherche Scientifique. LS was supported by a PhD Fellowship from La Ligue Contre le Cancer. We gratefully acknowledge the Image Analysis Hub and Bioinformatics and Biostatistics Hub (Research and Resource Centre for Scientific Informatics, Institut Pasteur) and the Fab Lab of Institut Pasteur for support in conducting this study. We gratefully acknowledge the UTechS Photonic BioImaging (Imagopole, C2RT, Institut Pasteur), supported by the French National Research Agency (France BioImaging, ANR-10-INBS-04; Investments for the Future), and acknowledge support from Institut Pasteur for the use of the spinning disk Nikon and the Trimscope multiphoton microscopes. We thank Christos Tsogkas, Wissal Manai and Benjamin Montagne for contributions to this study, and Dr. April Pyle for providing Seurat objects for scRNA-seq analysis^7^.

## Author contributions

Conceptualisation: LS, ST, BE; Methodology: LS, GL, BE; Software: LS, GL, HV, VL; Validation: LS, BE; Formal analysis: GL, HV, VL; Investigation: LS, JF, BE; Resources: ST; Data Curation: LS, GL, HV; Writing – original draft preparation: LS, ST, BE; Writing – review and editing: LS, GL, HV, JF, ST, BE; Visualisation: LS, GL, HV, BE; Supervision: ST, BE; Project administration: LS, ST, BE; Funding acquisition: ST, BE.

## Code availability

The code that was used in this study will be available on GitHub upon publication.

---

**Figure S1.**
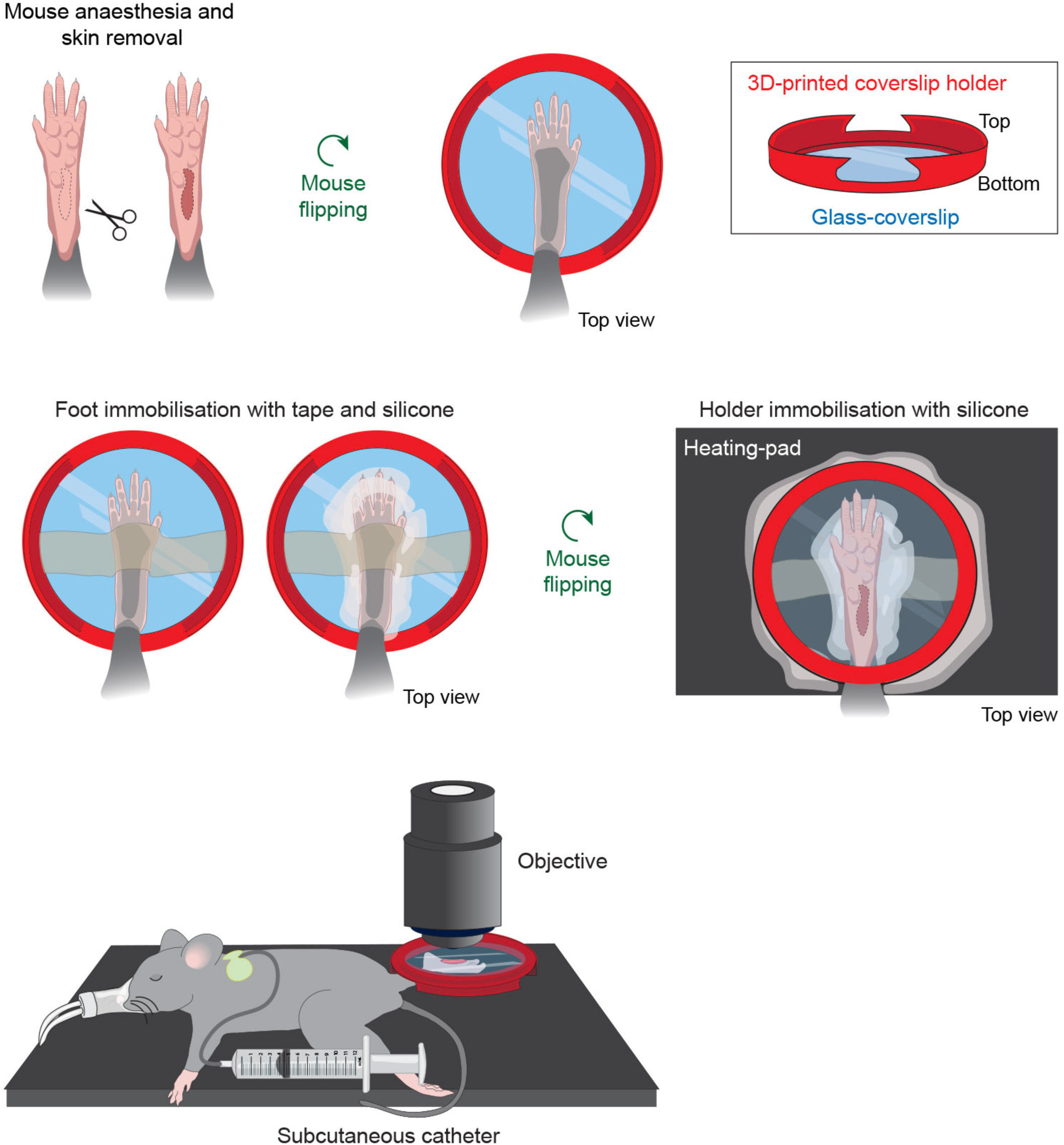
Intravital imaging setup. Related to Figure 1. Once anesthetised, a small piece of the mouse skin on foot was removed to expose the FDB. The foot was immobilised with tape and silicone to a glass-coverslip attached to a 3D-printed custom coverslip holder. The holder was fixed to a heating-pad (silicone) and the mouse was hydrated with 0.9% NaCl *via* a catheter, enabling continuous intravital imaging up to 10h. See also Methods section.

**Figure S2.**
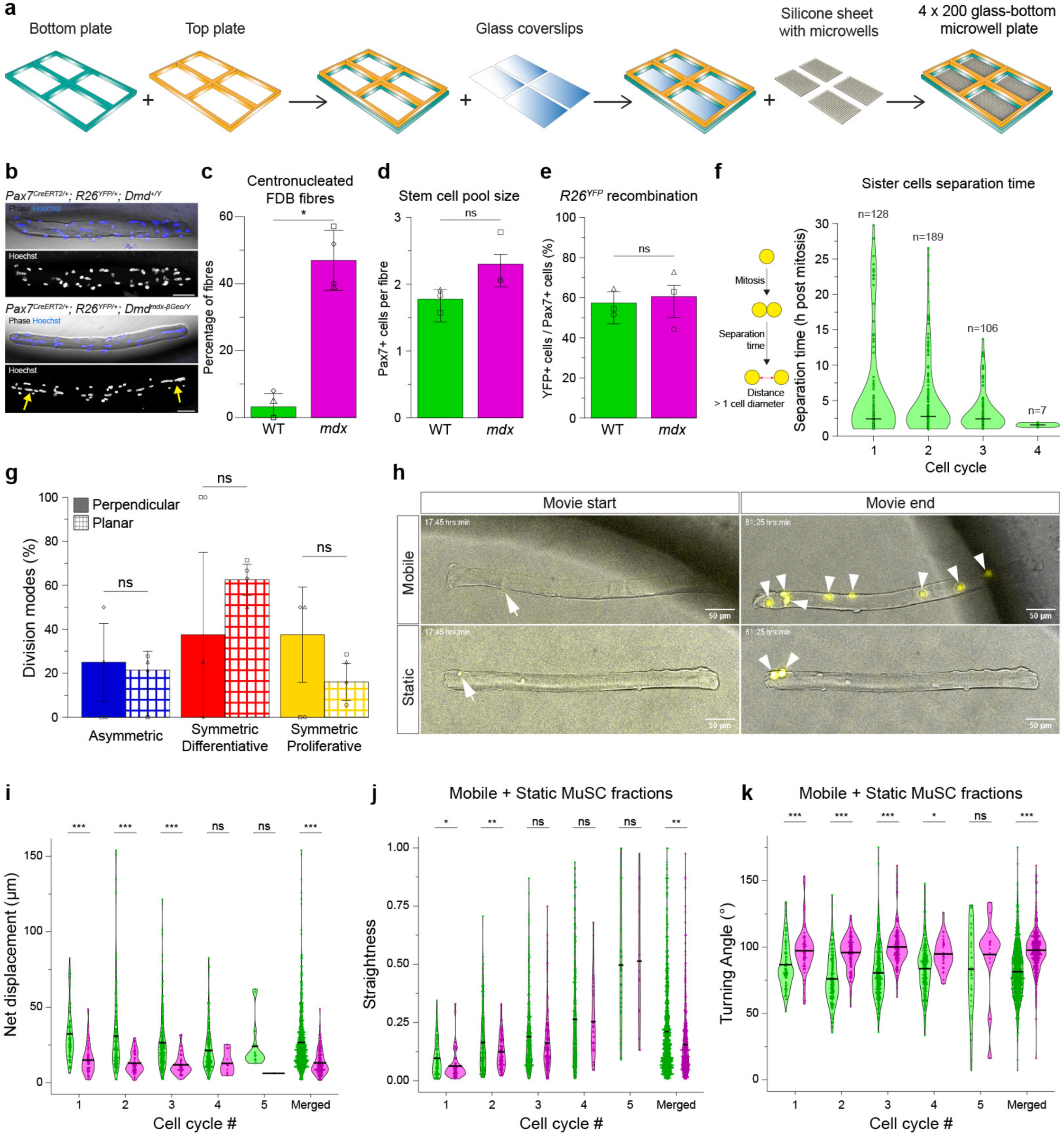
Related to Figure 2. **(a)** Microwell plate assembly. The microwell plate is composed of two plates (bottom and top) in which 4 rectangles were laser-cut, attached together. 4 glass coverslips were sealed to the outer part of the bottom plate and silicone sheets with microwells were attached on the glass coverslips. Single FDB myofibres were manually isolated, cultured, filmed and stained in each microwell. **(b)** Immunostaining of FDB fibres from *Pax7^CreERT2/+^; R26^YFP/+^*; WT and *mdx* 4-month-old mice. Phase contrast shows fibre shape; myonuclei labelled with Hoechst. Arrows indicate myonuclei in central position. Scale bar, 50 μm. **(c)** Percentage of centronucleated FDB fibres from *Pax7^CreERT2/+^; R26^YFP/+^*; WT and *mdx* 4-month-old mice. N = 4 experiments. p = 0.029. **(d)** Number of PAX7 positive MuSCs per FDB fibre from *Pax7^CreERT2/+^; R26^YFP/+^*; WT and *mdx* 4-month-old mice. Analysis performed immediately upon isolation. N = 3 experiments. p = 0.055. **(e)** Percentage of YFP-positive among PAX7-positive cells from FDB fibres of *Pax7^CreERT2/+^; R26^YFP/+^*; WT and *mdx* 4-month-old mice. Analysis performed immediately upon isolation. N = 3 experiments. p = 0.88. **(f)** Timing of sister cell separation (cell-cell distance larger than a cell diameter) after mitosis, at each individual cell cycles. N = 4 experiments, n = number of analysed divisions. **(g)** Modes of cell divisions of YFP-labelled myogenic *mdx* cells following perpendicular or planar divisions. N = 4 experiments. p-values: ACD (planar vs perpendicular) = 0.82; SCDd (planar vs perpendicular) = 0.19; SCDp (planar vs perpendicular) = 0.16. **(h)** Representative example of mobile (top) and static (bottom) migration of YFP-labelled myogenic cells (see Movie S4). **(i)** Net distance of mobile fraction of WT and *mdx* YFP-labelled myogenic cells, for individual and merged cell cycles. N = 4 experiments. **(j)** Straightness of mobile and static fractions of WT and *mdx* YFP-labelled myogenic cells, for individual and merged cell cycles. N = 4 experiments. **(k)** Turning angle of mobile and static fractions of WT and *mdx* YFP-labelled myogenic cells, for individual and merged cell cycles. N = 4 experiments. Statistical tests: **(c-e)** Wilcoxon test; **(g, i-k)** Linear mixed models. Horizontal lines in violin plots represent the mean. * *p* < 0.05, ** *p* < 0.01, *** *p* < 0.001.

**Figure S3.**
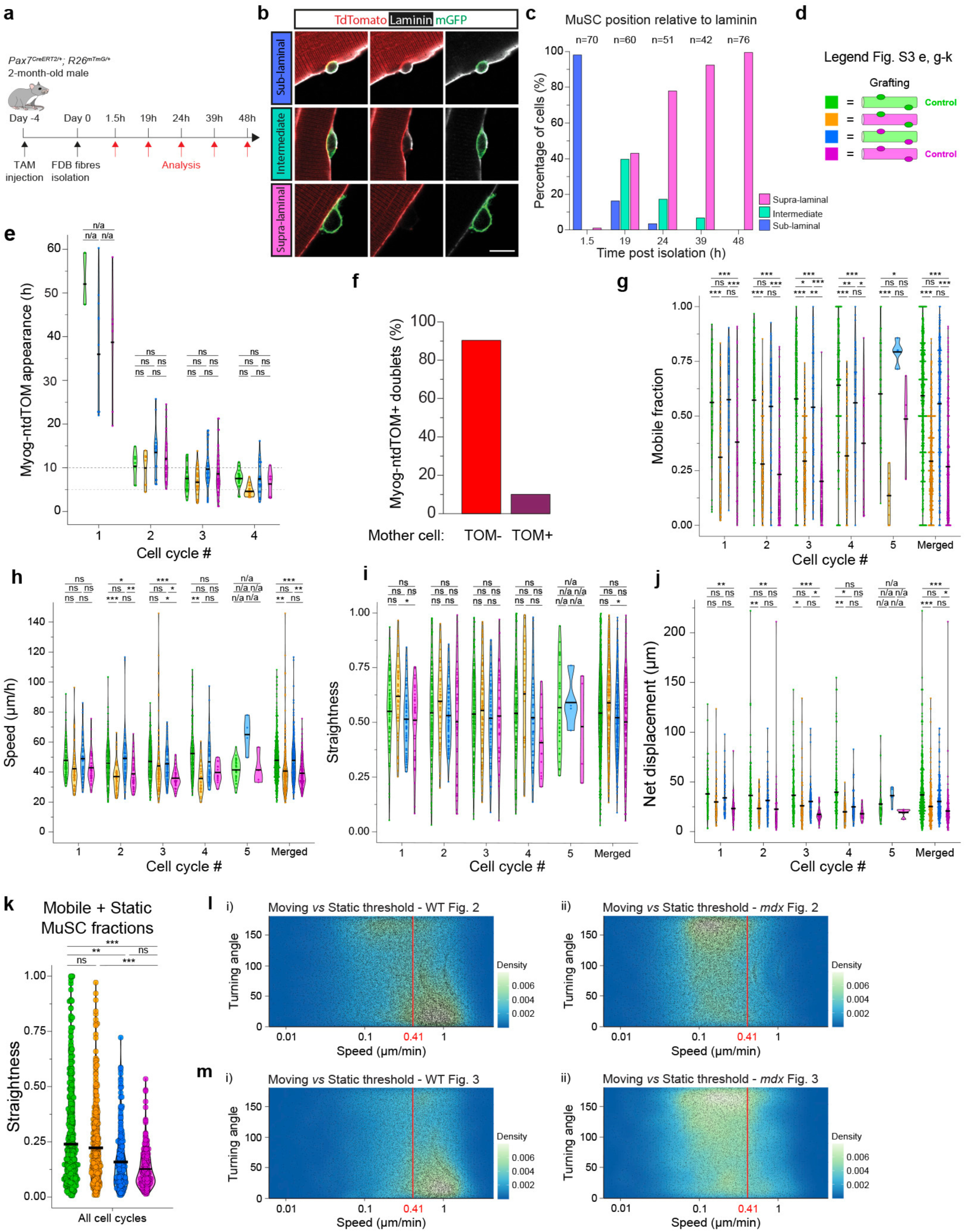
Related to Figure 3. **(a)** Experimental scheme. *Pax7^CreERT2/+^*; *R26^mTmG/+^* WT 2-month-old male mice were treated with tamoxifen to label MuSCs with GFP. FDB fibres were isolated, cultured, fixed and stained for laminin (basal lamina) at indicated time points post-isolation. **(b)** Representative example of sub-/supra-laminal or intermediate MuSC positions relative to basal lamina. **(c)** Percentage of sub-/supra-laminal or intermediate MuSC positions relative to basal lamina at different timepoints post-isolation. N = 3 experiment, n = analysed number of MuSCs. **(d)** Colour code of 4 grafting conditions (WT Ctrl, WT_to_*mdx, mdx*_to_WT, *mdx* Ctrl). **(e)** Timing of Myog-ntdTOM reporter appearance for each cell cycle, for all grafting conditions. N = 5 experiments. **(f)** Percentage of SCDd generating TOM+ doublet coming from a Myog-ntdTOM-positive or -negative mother cell. N = 5 experiments. **(g)** Mobile fraction for all grafting conditions for individual and merged cell cycles. N = 5 experiments. **(h)** Migration speed of mobile fraction for all grafting conditions for individual and merged cell cycles. N = 5 experiments. **(i)** Migration straightness of mobile fraction for all grafting conditions for individual and merged cell cycles. N = 5 experiments. **(j)** Net displacement of mobile fraction for all grafting conditions, for individual and merged cell cycles. N = 5 experiments. **(k)** Migration straightness of mobile and static fractions for all grafting conditions, for merged cell cycles. N = 5 experiments. **(l)** Threshold value between mobile and static fractions (Fig. 2 dataset). Density plots of turning angle over instantaneous speed for endogenous i) WT and ii) *mdx* MuSCs. **(m)** Threshold value between mobile and static fractions (Fig. 3 dataset). Density plots of turning angle over instantaneous speed for control grafting conditions i) WT Ctrl and ii) *mdx* Ctrl. Statistical tests: **(e, g-k)** Linear mixed models. Horizontal lines in violin plot represent mean. * *p* < 0.05, ** *p* < 0.01, *** *p* < 0.001.

**Figure S4.**
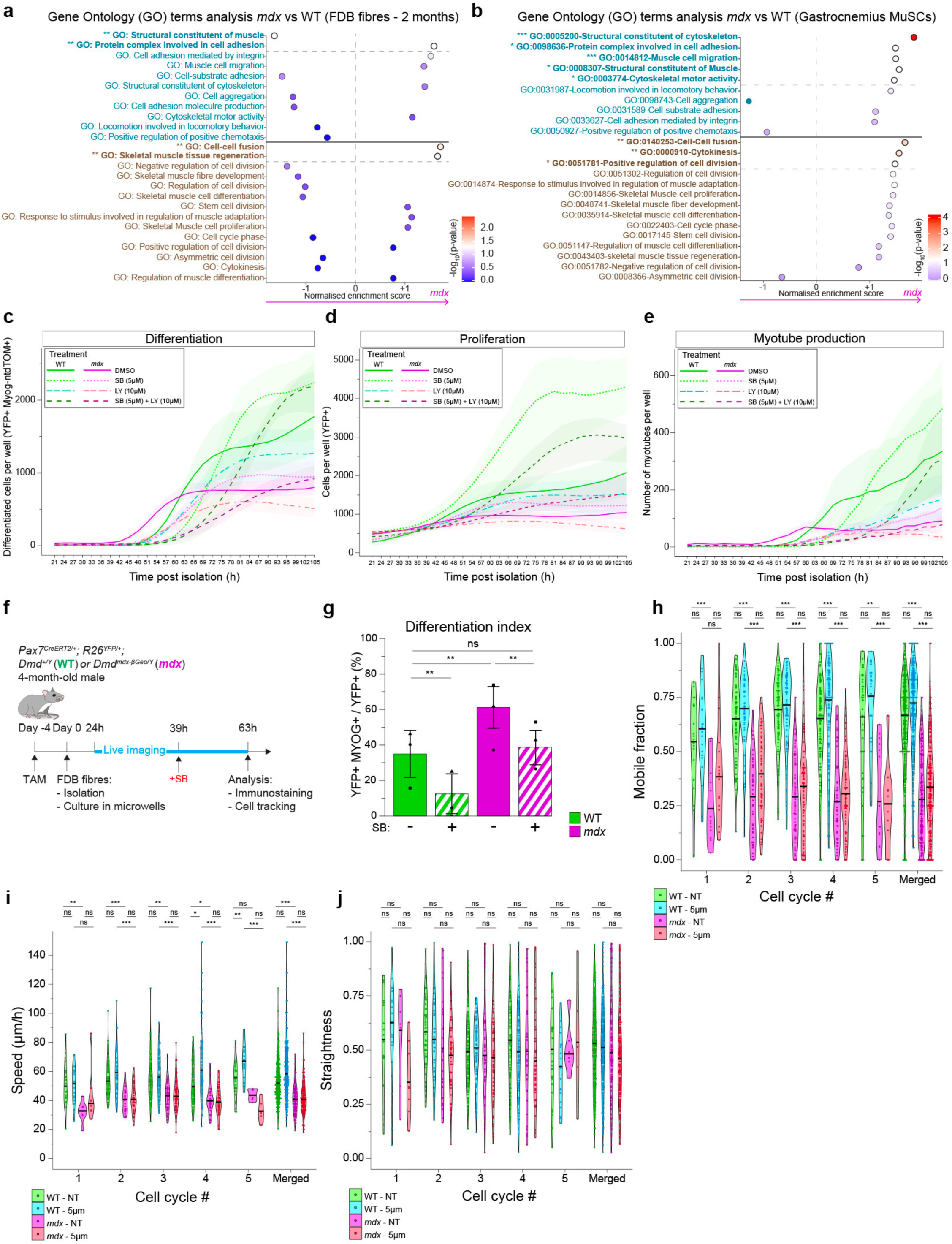
Related to Figure 4. **(a)** Gene Ontology (GO) enrichment analysis of FDB fibres RNA-seq data from 2-month-old WT and *mdx* mice^43^. Analysis focused on GO terms related to migration and proliferation/differentiation. **(b)** Gene Ontology (GO) enrichment analysis of gastrocnemius muscle scRNA-seq data from 2-month-old WT and *mdx* mice^7^. Analysis focused on GO terms related to migration and proliferation/differentiation in MuSC cluster. **(c)** Kinetics of differentiated cell production (number of YFP-positive Myog-ntdTOM-positive cells/well) of WT and *mdx* cells treated with SB and/or LY. N = 3 experiments. **(d)** Kinetics of total cell production (number of YFP-positive cells / well) of WT and *mdx* cells treated with SB and/or LY. N = 3 experiments. **(e)** Kinetics of myotube production (number of myotubes/well) of WT and *mdx* cells treated with SB and/or LY. N = 3 experiments. **(f)** Experimental scheme. *Pax7^CreERT2/+^; R26^YFP/+^; Dmd^+/Y^* (WT) and *Dmd^mdx-βGeo/Y^* (*mdx)* adult mice were treated once with tamoxifen. FDB fibres and associated MuSCs were isolated, cultured and filmed in microwells for ∼ 63h, fixed and immunostained. p38 inhibitor SB or vehicle (DMSO) were added 39h after isolation. **(g)** Differentiation index of WT and *mdx* YFP-labelled myogenic cells (∼ 63h of culture) with or without SB. N = 3 experiments. p (WT_Ctr vs *mdx*_Ctr) = 2.8e-3; p (WT_Ctr vs WT_SB) = 8.6e-3; p (*mdx*_Ctr vs *mdx*_SB) = 8.4e-3; p (WT_Ctr vs *mdx*_SB) = 0.42. N = 3 experiments. **(h)** Mobile fraction of WT and *mdx* YFP-labelled myogenic cells on isolated FDB fibres, with or without SB, for individual and merged cell cycles. N = 3 experiments. **(i)** Migration speed of mobile fraction of WT and *mdx* YFP-labelled myogenic cells on isolated FDB fibres, with or without SB, for individual and merged cell cycles. N = 3 experiments. **(j)** Migration straightness of mobile fraction of WT and *mdx* YFP-labelled myogenic cells on isolated FDB fibres, with or without SB, for individual and merged cell cycles. N = 3 experiments. Statistical tests: **(g)**Wilcoxon test; **(h-j)** Linear mixed models. Horizontal lines in violin plots represent the mean. * *p* < 0.05, ** *p* < 0.01, *** *p* < 0.001.

**Figure S5.**
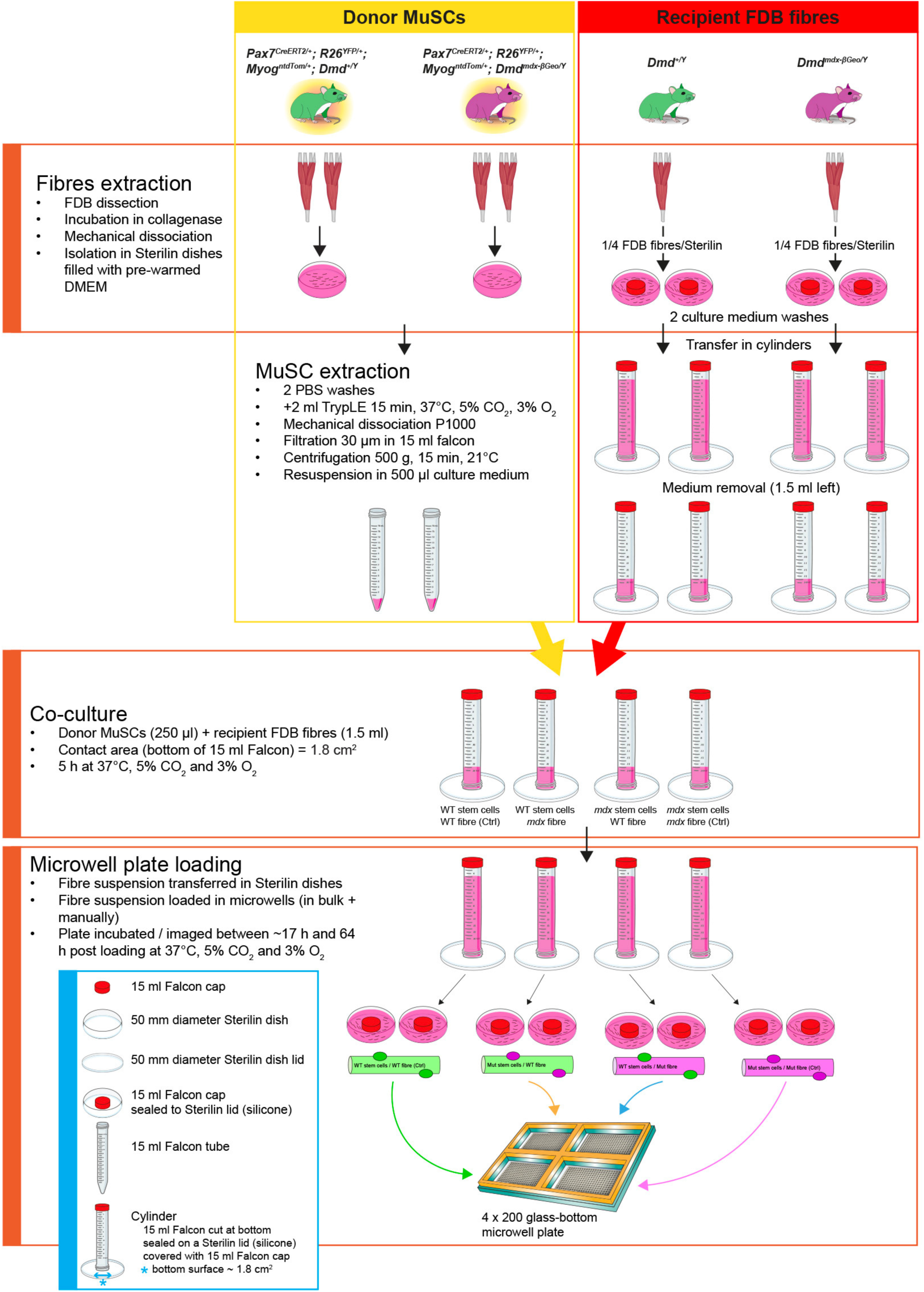
Protocol for cross-transplantation of MuSCs and FDB fibres. Related to Figure 3. Two *Pax7^CreERT2/+^; R26^YFP/+^; Myog^ntdTom/+^; Dmd^+/Y^* or *Dmd^mdx-βGeo/Y^* 4-month-old male mice were used as donors of WT or *mdx* YFP-labelled MuSCs respectively. Two *Dmd^+/Y^* or *Dmd^mdx-βGeo/Y^* 4-month-old male mice were used to generate WT or *mdx* recipient FDB fibres. FDB muscles from all mice were dissected and FDB fibres were isolated (see Methods). FDB fibres from donor mice were further treated (TrypLE, mechanical dissociation, filtration) to isolate MuSCs. Recipient FDB fibres mice were transferred in 1.5 ml of medium in cylinders (from Falcon 15 ml tubes). 4 combinations of donor MuSCs with recipient FDB fibres were co-cultured for 5h at 37°C, 5% CO_2_, 3% O_2_. Grafted FDB fibres were isolated in a microwell plate, cultured and live-imaged.

**Table S1. PCR primers**

**Table S2. Reagents**

**Movie S1.** Intravital imaging of injured FDBs at 3 dpi of *Pax7^CreERT2/+^; R26^mTmG/+^; Dmd^+/Y^* (WT, top) and *Dmd^mdx-βGeo/Y^* (*mdx,* bottom) adult mice. Related to Figure 1.

**Movie S2.** Intravital imaging of injured FDBs at 3 dpi of *Pax7^CreERT2/+^; R26^mTmG/+^; Dmd^+/Y^* (WT, left) and *Dmd^mdx-βGeo/Y^* (*mdx*, right) adult mice. High magnification of Movie S1 showing migrating myoblasts. Related to Figure 1.

**Movie S3.** Live imaging of FDB fibres and associated MuSCs in microwells from an adult *Pax7^CreERT2/+^; R26^mTmG/+^* mouse. Related to Figure 2.

**Movie S4.** Representative migration of mobile (top) and static (bottom) YFP-labelled myogenic cells on FDB fibres from a *Pax7^CreERT2/+^; R26^YFP/+^; Myog^ntdTom/+^; Dmd^+/Y^* mouse. Related to Figure 2.

**Movie S5.** Representative example of live imaging of a WT FDB fibre grafted with an *mdx* YFP; Myog-ntdTOM-labelled MuSC. Related to Figure 3.

**Movie S6.** Live-imaging of MuSCs from hindlimbs of *Pax7^CreERT2/+^; R26^YFP/+^; Myog^ntdTom/+^; Dmd^+/Y^* (WT, left) and *Dmd^mdx-βGeo/Y^* (*mdx*, right) mice isolated and cultured in 96 well plate. Related to Figure 4.

