## Supplementary material for "Impaired stem cell migration and divisions in Duchenne Muscular Dystrophy revealed by live imaging": Table S1

Table S1 - PCR primers

| Strain | Oligonucleotide | Sequence (5' to 3') | PCR conditions | PCR products |
|---|---|---|---|---|
| *Pax7* $^{CreERT2}$ | Pax7 GaKa mutant rev<br>Pax7 GaKa WT for<br>Pax7 GaKa WT rev | CAAAAGACGGCAATATGGTG<br>GCTGCTGTTGATTACCTGGC<br>CTGCACTGAGACAGGACCG | 95°C 5'<br>(95°C 30''; 55°C 30''; 72°C 1') x40<br>72°C 5' | WT: 419 bp<br>Knock-in: 235 bp |
| *R26* $^{mTmG}$ | CT2 R1 Rosa 1 WT<br>CT2 R3 Rosa 3 WT<br>pCAG R10 | AAAGTCGCTCTGAGTTGTTAT<br>GGAGCGGGAGAAATGGATATG<br>GTCGTTGGGCGGTCAG | 95°C 5'<br>(95°C 30''; 55°C 30''; 72°C 1') x40<br>72°C 5' | WT: 600 bp<br>Knock-in: 350 bp |
| *R26* $^{YFP}$ | CT2 R1 Rosa 1 WT<br>GFP seq2 for<br>Rosa26 F2 | AAAGTCGCTCTGAGTTGTTAT<br>CCGCCCTGAGCAAAGACCCCAACG<br>CAGGTTAGCCTTTAAGCCTGC | 95°C 5'<br>(95°C 30''; 55°C 30''; 72°C 1') x40<br>72°C 5' | WT: 247 bp<br>Knock-in: 392 bp |
| *Myog* $^{ntdTomato}$ | MyoG-nTdT Fwd<br>MyoG-nTdT Rev<br>MyoG-nTdT WT | TTCCTGTACGGCATGGACGAG<br>CAGGACAGCCCCACTTAAAAGC<br>CTTGCTGACCTGAGGGCC | 94°C 2'<br>(94°C 30''; 60°C 30''; 72°C 2') x34<br>72°C 10' | WT: 600 bp<br>Knock-in: 236 bp |
| *Dmd* $^{mdx\text{-}\beta Geo}$ | MDX intron 63R<br>MDX intron 63L<br>MDX 64<br>K03 | GCACGAGCATATGGTTGACACC<br>TAAGTTGAAAAGGTGAGGGC<br>CTCGCGGTTGAGGACAAACTCTTCGC<br>CGCATCGTAACCGTGCATCTGCCAGTTTGA | 95°C 5'<br>(95°C 30''; 55°C 30''; 72°C 1') x40<br>72°C 5' | WT: 200 bp<br>Knock-out: 350 bp |
