## Supplementary material for "Impaired stem cell migration and divisions in Duchenne Muscular Dystrophy revealed by live imaging": Table S2

Table S2 - Key Resources

| Reagent or Resource | Concentration | Supplier | Reference |
|---|---|---|---|
| **Antibodies** | | | |
| **Primary antibodies (Immunocytochemistry, ICC)** | | | |
| Mouse monoclonal anti-PAX7 | 1:20 | DSHB | Cat# PAX7 |
| Chicken polyclonal anti-GFP | 1:1000 | Abcam | Cat# ab13970 |
| Mouse monoclonal anti-MYOGENIN | 1:200 | DSHB | Cat# F5D |
| Rabbit polyclonal anti-LAMININ | 1:200 | Sigma | Cat# L9393 |
| **Secondary Antibodies (ICC)** | | | |
| Alexa Fluor 555 F(ab') Goat-anti-Mouse IgG1 | 1:500 | ThermoFisher | Cat# **A-21127** |
| Alexa Fluor 488 F(ab') Goat-anti-Chicken | 1:500 | ThermoFisher | Cat# **A-11039** |
| Alexa Fluor 633 Goat-anti-Rabbit | 1:500 | ThermoFisher | Cat# **A-21070** |
| **Chemicals, Peptides, and Recombinant Proteins** | | | |
| Imalgene 1000® | N/A | XX | XX |
| Rompun 2%® | N/A | XX | XX |
| Cardiotoxin | N/A | Latoxan | Cat# L8102 |
| Tamoxifen | N/A | Sigma | Cat# T5648 |
| Silicone | N/A | Smooth-on | Cat# MoldStar 20T |
| Ethanol 70% | N/A | Sigma | Cat# 32221 |
| Collagenase type 1 | N/A | Sigma | Cat# C0130 |
| Collagenase type 2 | N/A | Serlabo | Car# WOLS04177 |
| DNase I | N/A | Roche | Cat# 11284932001 |
| Ham's F10 | N/A | Sigma | Cat# N6635-10X1L |
| Dispase | N/A | Gibco | Cat# 17105-041 |
| DMEM GlutaMAX | N/A | ThermoFisher | Cat# 31966 |
| Horse Serum | N/A | ThermoFisher | Cat# 11510516 |
| Penicillin/Streptomycin | N/A | ThermoFisher | Cat# 15140122 |
| Chicken Embryo Extract | N/A | Life Science Production | Cat# MD-004D-UK |
| F12 | N/A | Fisher | Cat# 31765027 |
| Fetal MGI: 7442679 Serum | N/A | Fisher | Cat# 10-437-028 |
| Recombinant murine FGF-basic | N/A | PeproTech | Cat# 450-33 |
| Paraformaldehyde | N/A | Euromedex | Cat# 15710 |
| Hepes | N/A | Sigma | Cat# 51558-50ML |
| Goat Serum | N/A | ThermoFisher | Cat# 11540526 |
| Hoechst 33342 | N/A | ThermoFisher | Cat# H1399 |
| Triton | N/A | Sigma | Cat# T8787-250ml |
| TrypLE Express Enzyme | N/A | ThermoFisher | Cat# 10718463 |
| p38 MAPK inhibitor - SB 203580 | N/A | Sigma | Cat# 559389-5MG |
| PI3K inhibitors - LY 294002 | N/A | Cell signaling | Cat# 9901S |

Table S2 - Key Resources

| Experimental Models: Organisms/Strains | | | |
|---|---|---|---|
| Mouse *R26* $^{mTmG}$ | N/A | (Muzumdar et al., 2007) | MGI:3716464 |
| Mouse *Pax7* $^{CreERT2}$ | N/A | (Murphy et al., 2011) | MGI:5141477 |
| Mouse Rosa$^{YFP}$ | N/A | (Srinivas et al., 2001) | MGI: J:80963 |
| Mouse Dmd$^{mdx\text{-}\beta Geo}$ | N/A | (Wertz & Füchtbauer, 1998) | MGI: 94909 |
| Mouse Myogenin$^{ntdTomato}$ | N/A | (Benavente-Diaz et al., 2021) | MGI:7442679 |
| Mouse B6D2F1J | N/A | Janvier Labs | Cat# B6D2F1/JRj |
| **Software and Algorithms** | | | |
| Trackmate | N/A | (Tinevez et al., 2017) | N/A |
| Imaris 7.2.1 | N/A | Bitplane | N/A |
| ImageJ | N/A | N/A | N/A |
